# Immunological liver-skin axis: a systemic Vγ4^+^ γδT17 cell response links psoriasis-like inflammation and MASLD

**DOI:** 10.64898/2025.12.22.695890

**Authors:** Janik Fleißner, Yamila Rocca, Lukas Freund, Thomas Ossner, Fabian Imdahl, Rémi Doucet Ladevèze, Florian Kurz, Andreas Geier, Matthias Goebeler, Georg Gasteiger, Andreas Kerstan

## Abstract

Metabolic dysfunction-associated steatotic liver disease (MASLD) and psoriasis frequently co-occur and share interleukin-17 (IL-17)-mediated inflammatory mechanisms. Using a murine model combining Western-diet-induced early MASLD with chronic imiquimod-triggered psoriasis-like dermatitis, we observed exacerbated skin inflammation with increased cellularity and systemic IL-17–driven immune activation. In mice with psoriasis-like skin inflammation, hepatic γδT cells shifted from IFN-γ-producing γδT1 toward IL-17-secreting γδT17 cells, accompanied by an acute-phase response and induction of hepatic lipogenic gene expression. Single-cell RNA sequencing revealed an expansion of Vγ4⁺ γδT17 cells engaged in IL-17 and TGF-β signaling, consistent with early fibrogenic responses. Together, these findings identify Vγ4⁺ γδT17 cells as a mechanistic link between psoriasis and MASLD, promoting systemic inflammation and early hepatic remodeling. Consistently, in a clinical cohort of patients with moderate-to-severe psoriasis and co-morbid MASLD, IL-17-targeted therapy improved hepatic steatosis, underscoring the translational relevance of this immunological axis.

## Introduction

Metabolic dysfunction-associated steatotic liver disease (MASLD, formerly non-alcoholic fatty liver disease, NAFLD) has a global prevalence of 30%^1^ and is the second leading cause of end-stage liver disease^2^. MASLD is diagnosed when hepatic steatosis of no other identifiable etiology coexists with at least one cardiometabolic risk factor^3^. It comprises a spectrum from simple steatosis to steatohepatitis (MASH), which can progress to fibrosis, cirrhosis, and hepatocellular carcinoma^4^. Importantly, it is almost twice as common in patients with psoriasis, a chronic inflammatory skin and joint disease^5^. Both MASLD^6–8^ and psoriasis^9,10^ are driven by interleukin-17 (IL-17) signaling. While multiple biologics targeting IL-17 are approved for the treatment of moderate-to-severe psoriasis^11–14^, the thyroid hormone receptor beta-selective agonist resmetirom^15^ is currently the only approved therapy for MASH. Patients with either condition are at increased risk of metabolic syndrome and cardiovascular disease^16,17^. Of note, weight loss has demonstrated clinical benefits in both patient groups^18–20^.

In mice, skin inflammation resembling psoriasis can be robustly induced by repeated topical application of the toll-like receptor 7 agonist imiquimod (imq)^21^. Reflecting the impact of nutrition and obesity on psoriatic skin inflammation, several studies have demonstrated that lipid and carbohydrate components of high-fat (HFD) and Western-type diets (WD) aggravate imq-induced psoriasis-like inflammation in mice^22–25^. Mechanistically, diet-derived oxysterols intensify inflammation by modulating the peripheral function of dermal IL-17-producing γδT (γδT17) cells^26^. Furthermore, oxidative stress from free-fatty-acid breakdown depletes dermal PPARγ⁺ regulatory T cells, which reduces the suppression of γδT17 cells and promotes inflammation^27^. Under psoriatic conditions, γδT17 cells metabolically reprogram toward increased glycolysis and acetyl-CoA carboxylase 1-dependent fatty acid synthesis, which enhances IL-17A production and promotes inflammation^28^. However, these studies predominantly investigated acute (single-phase) psoriasis-like dermatitis models, whereas patients typically experience chronic, relapsing-remitting disease.

γδT cells are part of the unconventional T cell compartment. They are enriched at barrier tissues, where they act as sentinels and orchestrate local immune responses^29^. In murine skin, γδT cells maintain homeostasis and contribute to wound healing^30^. According to the Heilig and Tonegawa nomenclature^31^, Vγ5^+^ dendritic epidermal γδT cells (DETCs)^32^ as well as Vγ4^+^ and Vγ6^+^ dermal γδT cells^33,34^ can be differentiated in murine skin. In the liver, Vγ1^+35^ and Vγ6^+36^ γδT cells predominate. While their precise role in liver diseases remains incompletely understood, hepatic γδT cells have been implicated in MASLD development, viral hepatitis^37,38^ and fibrogenesis^39^. In a dietary model of MASLD, γδT17 cells accelerated disease progression, whereas *Tcrd^-/-^*mice, which lack γδT cells, were protected from liver damage^40^. In models of MASLD and alcohol-induced steatohepatitis, γδT cells were recruited to the liver, adopted a γδT17 phenotype, and sustained inflammation by suppressing protective IFN-γ production in CD4⁺ T cells^41^.

Given the involvement of γδT17 cells in severe MASLD and acute psoriasis-like inflammation, and the limited availability of models reflecting the relapsing nature of human psoriasis, we established a model of early diet-induced MASLD combined with repeated imiquimod-triggered psoriatic flares to assess whether recurrent skin inflammation promotes hepatic disease progression via systemic Vγ4⁺ γδT17 cells. Our study demonstrates a γδT17 cell-driven skin-liver crosstalk linking psoriasis and MASLD. This murine immunological landscape may parallel the human context, with analogous γδT cell subsets, additional IL-17-producing immune populations, and converging inflammatory pathways. Consistently, in our clinical cohort of patients with moderate-to-severe psoriasis and co-morbid MASLD, IL-17-targeted systemic therapy resulted in an improvement of hepatic steatosis, highlighting the translational relevance of this immunological axis.

## Results

To investigate whether recurrent psoriasis-like skin inflammation influences hepatic pathology in the context of diet-induced MASLD, a question which has not been addressed in previous acute imiquimod models, we used mice fed either a chow diet or a WD enriched with fructose for 16 weeks to induce early MASLD (Fig. S1A). To mimic the relapsing inflammatory nature of human psoriasis, mice underwent three cycles of topical imiquimod (imq) treatment, while control animals received vaseline (mock) (Fig. S1B).

### Western-type diet induces early MASLD characteristics

Mock-treated mice fed a WD (mock WD) showed a significantly higher weight gain over the course of the diet compared to their chow-fed controls (mock chow) (Fig. 1A). At the end of the diet, the mean body weight was 15.3 % higher in comparison to the chow controls (28.0 ± 0.7 g (SEM) vs. 24.3 ± 0.7 g). At sacrifice, the mean liver weight of mock WD was 49.2 % higher than that of mock chow (1.95 ± 0.06 g vs. 1.31 g ± 0.04 g), and the mean liver-to-body-weight ratio was significantly elevated (7.07 ± 0.19 % vs. 5.34 %, ± 0.15 %) (Fig. 1B). Histologically, mock WD livers exhibited increased steatosis (21.4 ± 3.9 % of area in quantification per high power field vs. 1.3 ± 0.3 %) (Fig. 1C). Although hepatic lipids reflected the histologically observed lipid accumulation only partly, mock WD livers displayed significantly increased amounts of cholesterol and a numerically higher triglyceride content (1.8-fold ± 0.2 and 2.1-fold ± 0.5, respectively) (Fig. 1D). Consistent with steatosis, hepatic mRNA expression of key enzymes regulating lipogenesis was significantly overexpressed (Fig. 1E). Compared to mock chow, the gene expression of *Mogat1,* encoding monoacylglycerol O-acyltransferase 1, was strongly induced (18.9-fold ± 2.4). In addition, *Cd36*, encoding fatty acid translocase, and *Scd1*, encoding stearoyl-CoA desaturase, were transcriptionally up-regulated (4.8-fold ± 0.5 and 4.5-fold ± 0.4, respectively). *Acaca*, encoding acetyl-CoA carboxylase, and *Fasn*, encoding fatty acid synthase, were not regulated by WD (1.3-fold ± 0.2 and 1.1-fold ± 0.2, respectively). Consistent with early MASLD, WD led to a significant induction of *Cxcl2* and *Ccl2* mRNA (3-fold ± 0.7 and 2.3-fold ± 0.3, respectively), suggesting recruitment of neutrophils and monocytes to the liver, while *Il6* was not transcriptionally regulated (1.2-fold ± 0.3) (Fig. 1F). In mock WD, alanine aminotransferase (ALT) plasma activity was significantly elevated (mock chow: 27.8 ± 1.7 U/L vs mock WD: 44.1 ± 3.2 U/L, *p* = 0.005) (Fig. 1G), demonstrating the onset of hepatocyte injury. Together, these findings confirm that WD induces early MASLD in our model.

**Figure 1.**
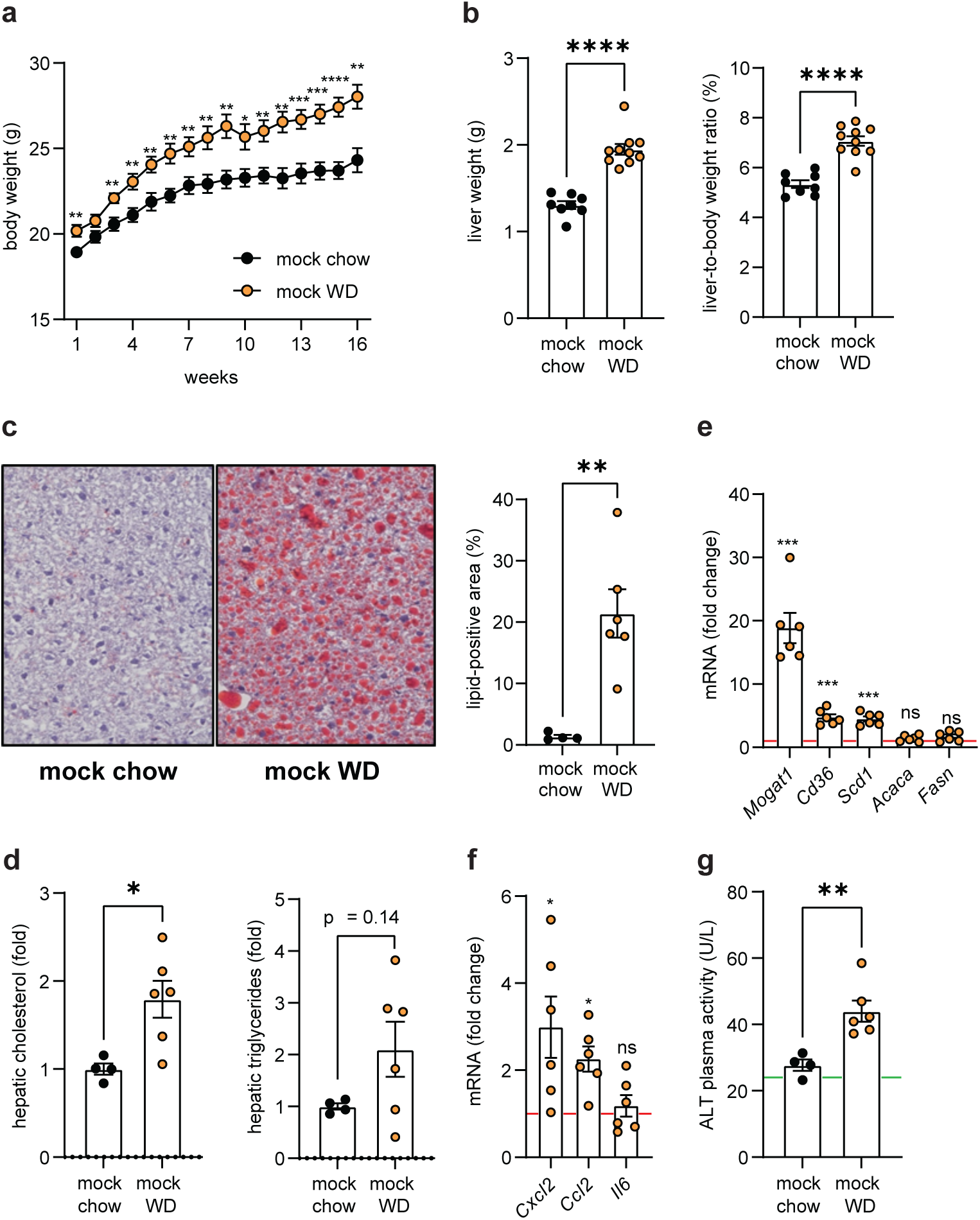
Western-type diet induces features of early MASLD. (**a**) Body weight (g) over the course of chow and WD. Data are pooled from two independent experiments (mock chow, n = 8; mock WD, n = 10). (**b**) Liver weight (left, g) and liver-to-body weight ratio (right, %) at the time of sacrifice (mock chow, n = 8; mock WD, n = 10). Data pooled from two independent experiments. (**c**) Representative Oil Red O-stained liver sections (left) and quantification of lipid-positive area per high-power field (right), (mock chow, n = 4; mock WD, n = 6). (**d)** Hepatic cholesterol (left) and triglyceride (right) content, determined by colorimetric assays following Folch lipid extraction, shown relative to mock chow (mock chow, n = 4; mock WD, n = 6). (**e, f**) Relative qPCR analyses of genes involved in hepatic lipid metabolism (E) as well as proinflammatory cytokine and chemokine genes (F). Results of mRNA expression were normalized to *Rplp0* and presented relative to mock chow (mock chow, n = 4; mock WD, n = 6). (**g**) Plasma activity of alanine aminotransferase (ALT, U/L) (mock chow, n = 4; imq WD, n = 6 each). The green solid line indicates the median level of ALT activity reported for healthy female mice aged 90–135 days^68^.

### Early MASLD aggravates imq-induced, γδT17-driven chronic psoriatic skin inflammation

Upon chronic imq treatment, WD-fed mice (imq WD) developed a more pronounced psoriatic skin inflammation than imq-treated chow-fed controls (imq chow) (Fig. 2A and C). We clinically scored the skin using the modified Psoriasis Area and Severity Index (mPASI), encompassing erythema, induration, and desquamation. Imq WD exhibited significantly higher total mPASI scores than all other groups, with both erythema and desquamation scores exceeding those of imq chow (Fig. 2C). Clinical response to imq treatment reflected by total mPASI did not differ between the initial exposure and the re-challenge (data not shown). This was paralleled by a significant increase in skin immune cellularity, as determined by flow cytometry quantification of live CD45^+^ cells (Fig. 2B). Of note, the number of cells was significantly higher in mock WD than in mock chow (9711 ± 1730 vs. 2451 ± 668 cells per 100 mg skin). In line with the histopathological features of human psoriasis, imq-treated mice exhibited profound keratinocyte hyperplasia (Fig 2D). Like the parameter ‘induration’ in the mPASI (Fig. 2B), this was not influenced by diet (imq WD: 104.4 ± 6.5 µm vs. imq chow: 90.0 ± 6.3 µm).

**Figure 2:**
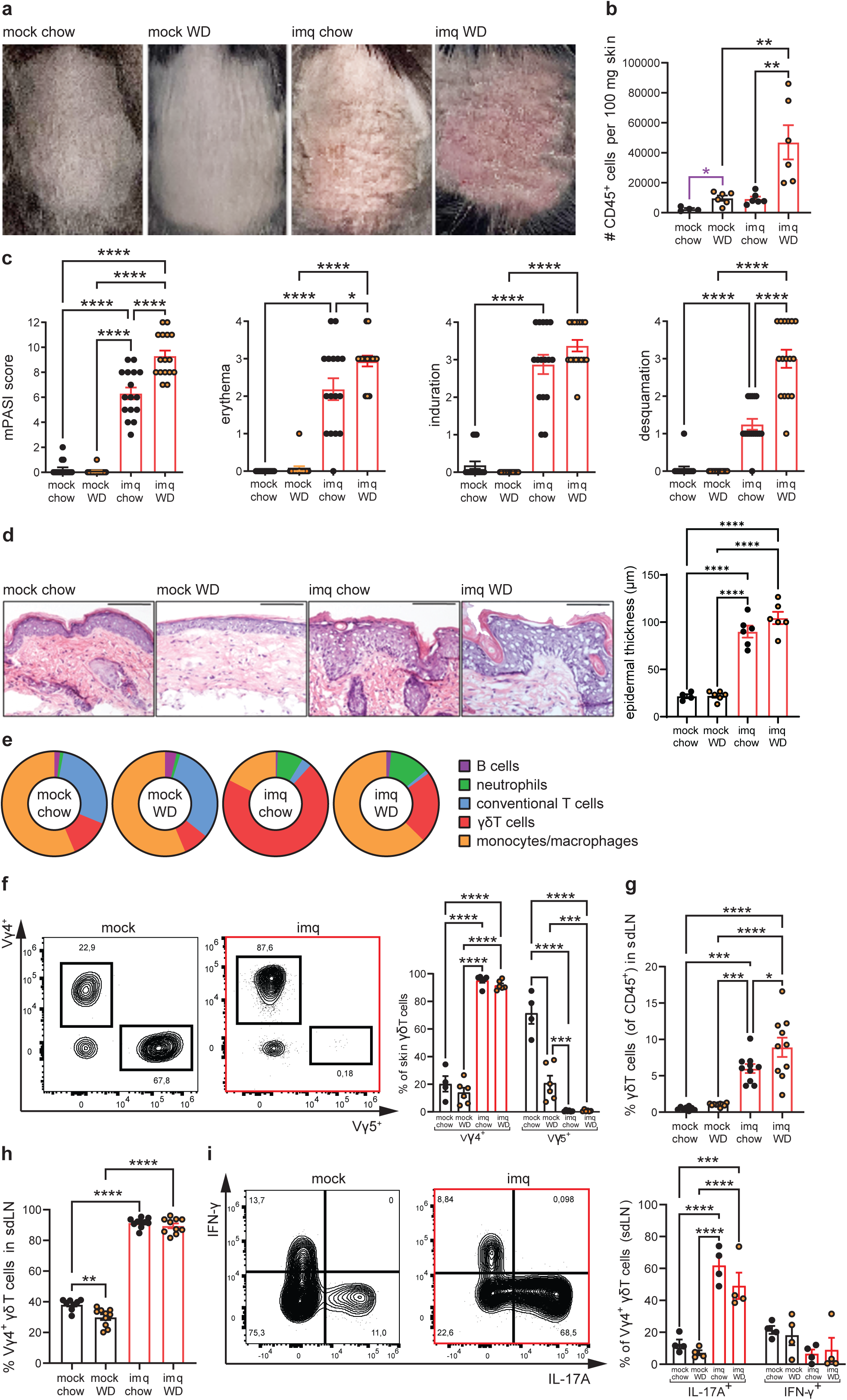
Early MASLD aggravates imq-induced, γδT17-driven chronic psoriatic skin mmation. (**a**) Representative clinical photographs of mouse skin obtained one day after the last topical treatment. (**b**) Absolute number of live CD45^+^ immune cells per 100 mg skin, quantified by flow cytometry (mock chow, n = 4; imq chow, mock WD, and imq WD, n = 6 each). (**c**) Modified Psoriasis Area and Severity Index (mPASI) considering erythema, induration, and desquamation with each category being scored 0 to 4, independently assessed by two dermatologists. Data pooled from 4 independent experiments (mock chow, imq chow, and imq WD, n = 16 each; mock WD, n = 15). (**d**) Representative H&E-stained skin sections (left, scale bar indicating 100 μm) and quantification of epidermal thickness (right) (mock chow, n = 4; imq chow, mock WD, and imq WD, n = 6 each). (**e**) Pie charts showing mean distribution of major skin-resident CD45^+^ immune subsets defined as: CD19^+^ (B cells), CD19- CD11b^+^Ly6G^+^ (neutrophils), CD19-Ly6G-CD11b-CD90.2^+^CD3^+^NKp46-TCRγδ^+^ (γδT cells), CD19-Ly6G-CD11b-D90.2^+^CD3^+^NKp46-TCRγδ- (conventional T cells), and CD19-Ly6GCD11b^+^ F4/80^+^ (monocytes/macrophages), (mock chow, n = 4; imq chow, mock WD, and imq WD, n = 6 each). (**f**) Frequencies of Vγ4^+^ and Vγ5^+^ γδT cells within skin γδT cells. Representative contour plots (left) of mock chow and imq chow and bar graphs (right), indicating mean percentages of (mock chow, n = 4; imq chow, mock WD, and imq WD, n = 6 each). (**g**) Proportion of γδT cells (CD19-Ly6G- CD11b-F4/80-CD90.2^+^CD3^+^TCRγδ^+^) within live CD45^+^ cells in skin-draining lymph nodes (sdLN) (mock chow, n = 8; imq chow, mock WD, and imq WD, n = 10 each). (**h**) Frequency of Vγ4^+^ γδT cells within γδT cells in sdLN (mock chow, n = 8; imq chow, mock WD, and imq WD, n = 10 each). (**I**) IL-17A and IFN-γ expression of Vγ4^+^ γδT cells in sdLN, gated on live CD45^+^CD19-CD90.2^+^CD3^+^F4/80-CD4- CD8-TCRγδ^+^TCRVγ4^+^. Representative contour plots (left) of mock chow and imq chow and quantification (right) (n = 4 per group). One-way ANOVA, Kruskal-Wallis test, or unpaired ttest (violet).

Next, we used flow cytometry to assess changes in the composition of skin immune cells (Fig. 2E). Besides higher percentages of neutrophils, we observed a pronounced increase in γδT cells accompanied by a reduction in conventional T cells in both imq chow and imq WD compared to mock. Given the crucial role of IL-17-producing γδT cells (γδT17 cells) in the imq model of psoriasis^42^, we focused on this subset of unconventional T cells. Independent of diet, imq treatment induced a shift from Vγ5^+^ DETCs to Vγ4^+^ dermal γδT cells (Fig. 2F). In skin-draining axillary and inguinal lymph nodes (sdLN) of imq-treated mice, we detected an increase in total γδT cells, with imq WD showing a significantly higher frequency than imq chow (Fig. 2G). The proportion of Vγ4^+^ cells was again markedly increased in sdLN of imq-treated mice (Fig. 2H), displaying a similarly activated CD44^hi^CD62L^lo^CD103^+^ phenotype as their skin-resident counterparts (Fig. S2B). In both skin and sdLN, imq treatment induced an increased RORγt expression and reduced CD27 expression in these cells (Fig. S2C). Upon PMA/ionomycin *ex vivo* stimulation, cytokine production of Vγ4⁺ γδT cells in sdLN shifted from an interferon-γ(IFN-γ)-favoring profile in mock mice to a dominant IL-17A response in imq-treated mice (Fig. 2I).

Thus, while MASLD-inducing WD aggravated psoriatic skin inflammation, imq treatment led to an increased frequency of Vγ4^+^ γδT17 cells in both skin and sdLN.

### Coexistence of early MASLD and chronic psoriasis synergistically contribute to systemic inflammation and induction of hepatic lipogenesis

During each cycle of imq treatment, mice displayed weight loss, which was only transient in imq chow (Fig. 3A). By contrast, imq WD exhibited a significantly greater weight loss than imq chow (16.0 ± 1.0 % vs. 2.5 ± 1.4), which persisted for approximately two weeks (Fig S3A). In line with aggravated disease, imq WD led to the numerically highest liver-to-body weight ratio (Fig. 3B). Relative liver weight was significantly increased in both imq-treated groups compared with chow-fed controls (imq chow: 8.0% vs. mock chow: 5.3%; imq WD: 8.6% vs. mock WD: 7.1%). Of note, livers of imq WD displayed a higher immune cell count than those of imq chow (207,215 vs. 148,556 cells per 100 mg) (Fig. 3C). In addition, imq-treated mice developed splenomegaly (Fig. S3B), a known feature of the imq model of psoriasis^21^, indicative of systemic inflammation. Consistently, plasma cytokine analysis (Fig. 3D) revealed an aggravated inflammatory profile in imq WD. IL-17A and IL-17F were most elevated in imq WD, exceeding mock controls and trending higher than imq chow (190.8 vs. 85.8 pg/mL and 6.6 vs. 2.9 pg/mL, respectively). IL-10 and tumor necrosis factor-α (TNF-α) were increased in imq WD compared to all other groups, while IL-6 showed modest increases in both imq-treated groups relative to mock WD, though absolute levels remained within the normal range. IFN-γ and IL-22 were unchanged and below reference levels (Fig. S3C). At the transcriptional level, hepatic expression of genes encoding hepatic acute-phase proteins (*Saa3, Saa1, Lcn2*) was significantly induced in imq WD, further corroborating a systemic inflammatory state (Fig. 3E). Notably, *Srebf1*, encoding sterol regulatory element-binding transcription factor 1 (SREBP1), a key regulator of fatty acid synthesis and lipogenesis, was significantly up-regulated in livers of imq WD compared with all other groups (3.2-fold ± 0.6 relative to mock chow, Fig. 3F).

**Figure 3:**
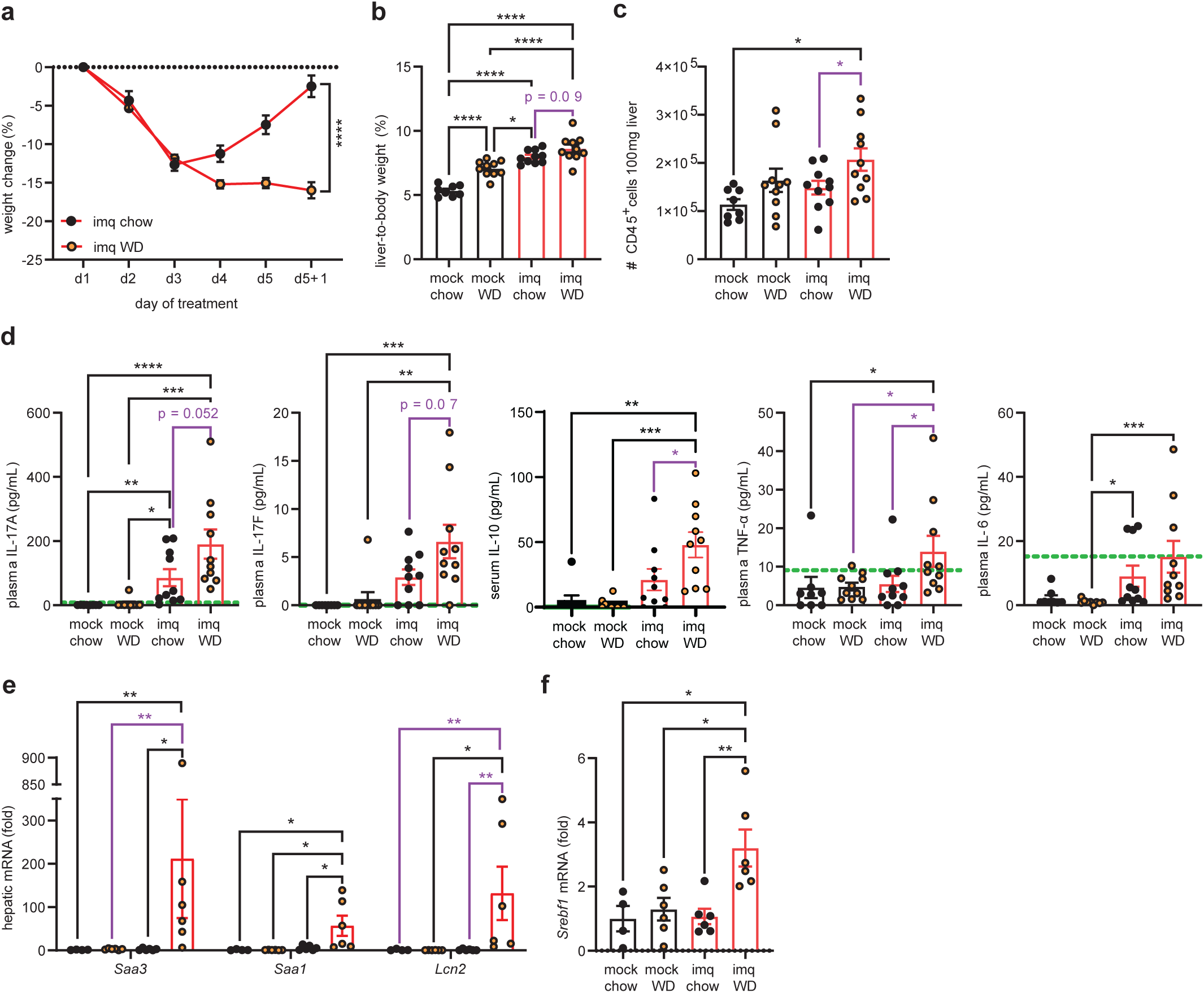
Coexistence of early MASLD and chronic psoriasis synergistically contribute to systemic inflammation and induction of hepatic lipogenesis. (**a**) Body weight change during each skin treatment cycles, presented as a percentage change relative to day (d1). Data pooled from 7 independent experiments (mock WD, *n* = 35, and imq WD, *n* = 36). (**b**) Liver-to-body weight ratio (%) at the time of sacrifice (mock chow, n = 8; imq chow, mock WD, and imq WD, n = 10 each). **(c)** Absolute number of live CD45^+^ immune cells per 100 mg liver, quantified by flow cytometry (mock chow, n = 8; imq chow, mock WD, and imq WD, n = 10 each). (**d**) Plasma concentrations of IL-17A, IL-17F, IL- 10, TNF-α, and IL-6 in pg/mL (mock chow, n = 8; imq chow, mock WD, and imq WD, n = 10 each). Duplicate measurements per cytokine were averaged. Green dashed lines indicate mean concentrations in healthy controls as reported by the manufacturer. (**e**) Relative qPCR analyses of genes involved in hepatic acute-phase response and (**f**) regulation of lipid metabolism. Results of mRNA expression were normalized to *Rplp0* and presented relative to mock chow (mock chow, n = 4; imq chow, mock WD, and imq WD, n = 6 each). One-way ANOVA or Kruskal-Wallis test for multiple comparisons; unpaired t-test or Mann-Whitney test for comparison of two groups (violet).

Together, these results demonstrate that chronic psoriatic inflammation in the context of early MASLD not only amplifies systemic inflammation but also drives hepatic lipogenesis.

### Early MASLD with chronic psoriatic skin inflammation enriches hepatic Vγ4⁺ γδT cells with an IL-17-skewed profile and loss of γδT1 dominance

Next, we analyzed hepatic immune cell composition by flow cytometry (Fig. 4A). Similar to the skin, imq treatment consistently reduced B and conventional T cell frequencies, while neutrophils, monocytes/macrophages, and γδT cells were increased. Livers of imq-treated mice showed significantly higher proportions of γδT cells than controls (imq chow: 2.6 ± 0.5% vs. mock chow: 0.6 ± 0.1; imq WD: 2.7% ± 0.2% vs. mock WD: 0.8 ± 0.1%) (Fig. 4B). Again, γδT cells in imq-treated mice were predominantly Vγ4^+^ (imq chow: 82.9 ± 2.1% vs. mock chow: 32.1 ± 2.0; imq WD: 78.5% ± 2.4% vs. mock WD: 28.0 ± 1.4%) (Fig. 4C). Of interest, these abundant RORγt^+^CD44^hi^CD27^-^ γδT17 cells outnumbered Eomes^+^CD44^hi^CD27^+^ γδT1 cells, which predominated in livers of mock-treated mice. (Fig. 4D). Functional characterization after *ex vivo* PMA/Ionomycin stimulation revealed a loss of IFN-γ-producing Vγ4^+^ γδT cells in livers of imq-treated mice (Fig. 4E). Interestingly, Vγ4^+^ γδT cells of imq WD livers displayed significantly higher basal expression of the activation marker CD69 than those of imq chow (16019 ± 1300 vs. 22460 ± 1467; MFI) (Fig. 4F). Upon stimulation, CD69 expression was robustly up-regulated in mock-treated mice but remained unchanged in imq chow and was significantly downregulated in imq WD relative to their respective unstimulated control. In contrast, Vγ4^+^ γδT cells from sdLN showed a strong induction of CD69 MFI upon stimulation across all groups (Fig. 4F). These results suggest that the coexistence of early MASLD and chronic psoriatic skin inflammation selectively favors Vγ4⁺ γδT17 cells in the liver at the expense of IFN-γ-producing γδT1 cells.

**Figure 4:**
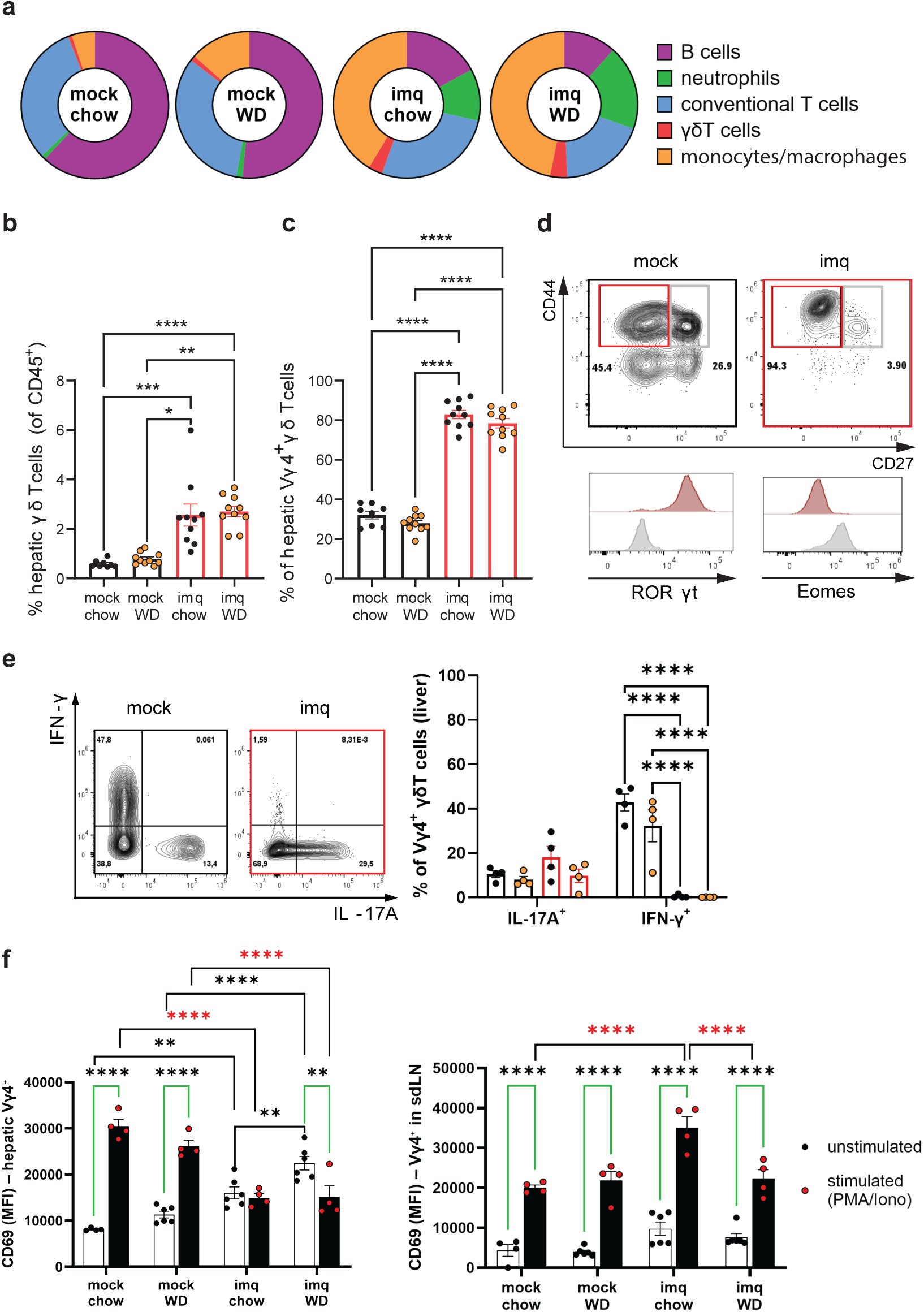
Early MASLD with chronic psoriasis enriches hepatic Vγ4⁺ γδT cells with an IL-17-skewed profile and loss of γδT1 dominance. (**a**) Pie charts showing mean distribution of major liver CD45^+^ immune subsets defined as: CD19^+^ (B cells), CD19-CD11b^+^Ly6G^+^ (neutrophils), CD19-Ly6G-CD11b-CD90.2^+^CD3^+^NKp46- TCRγδ^+^ (γδT cells), CD19-Ly6G-CD11b-CD90.2^+^CD3^+^NKp46-TCRγδ- (conventional T cells), and CD19-Ly6G-CD11b^+^F4/80^+^ (monocytes/macrophages), (mock chow, n = 4; imq chow, mock WD, and imq WD, n = 6 each). (**b**) Frequency of γδT cells per 100 mg liver tissue, shown as percentage of live CD45⁺ cells quantified by flow cytometry (mock chow, n = 8; imq chow, mock WD, and imq WD, n = 10 each). (**c**) Frequency of Vγ4⁺ γδT cells within hepatic γδT cells. (**d**) Representative contour plots of CD27 and CD44 expression in hepatic Vγ4⁺ γδT cells from mock chow and imq chow (top). Mean fluorescence intensity (MFI) of RORγt and Eomes in CD44hiCD27⁺ (grey) and CD44hiCD27⁻ (dark red) hepatic Vγ4⁺ γδT cells from a representative imq chow sample (bottom). (**e**) IL-17A and IFN-γ expression of hepatic Vγ4^+^ γδT cells. Representative contour plots (left) and quantification (right) of mock chow and imq chow and (n = 4 per group). (**f**) Vγ4^+^ γδT cells from liver (left) and sdLN (right) were left untreated or stimulated with PMA/ionomycin, and CD69 expression was analyzed by flow cytometry. Green brackets indicate comparisons between stimulated and unstimulated conditions; red asterisks indicate differences within stimulated groups.

### Single-cell RNA sequencing identifies hepatic γδT17 cells engaged in TGF-β signaling in early MASLD with concurrent chronic psoriasis

To study hepatic immune cell subsets in the context of chronic psoriasis and early MASLD, we performed single-cell RNA sequencing (scRNA-seq). Unsupervised clustering followed by UMAP projection revealed 13 transcriptionally distinct clusters (Fig. 5A). Annotation based on canonical lineage-defining genes identified one cluster of T helper cells (cluster 0: *Cd3d*, *Cd4),* three clusters of cytotoxic T cells (clusters 1, 4, and 5: *Cd3d*, *Cd8*), three clusters of monocytes/macrophages (clusters 2, 6, and 11: *Cd14*, *Adgre1*), two clusters of γδT cells (clusters 3 and 10: *Trgv1*, *Trgv2*, *Trgc2*), one cluster of ILC1/NKT cells (cluster 7: *Klrb1, Ncr1, Cd3d*), one cluster of NK cells (cluster 8: *Eomes, Klrb1, Ncr1*), one cluster of neutrophils (cluster 9: *Ly6G, Ly6c2*), and one cluster of eosinophils/basophils (cluster 12: *Mcpt8, Siglecf*) (Fig. 5B). Principal component analysis (PCA) and UMAP projection demonstrated that transcriptional profiles aligned with imq treatment rather than diet (Fig. 5C and Fig. S4). ScRNA-seq data were aggregated by experimental condition to generate pseudobulk expression profiles, which were subsequently analyzed using the *RunPCA* function. Under mock conditions, samples differed according to diet (WD vs. chow), a separation that was primarily captured along the PC2 axis. Upon imq treatment, however, the dietary effect became less pronounced, and variation along the PC1 axis instead reflected the distinction between imq-treated and untreated samples. Nevertheless, cluster distribution showed that circulating central memory CD8^+^ T cells (Tcm, cluster 1: *Sell*, *Ccr7*, *Il7r*; Fig. S5A) were less abundant in livers of WD-fed mice, whereas NK cells were enriched (Fig. 5D). Consistent with our flow cytometry data, proportions of γδT cells and neutrophils were increased in livers of imq-treated mice, while the frequency of a monocyte/macrophage population (cluster 6) was higher in mock-treated mice. Of note, cycling CD8^+^ T cells (clusters 4 and 5) and γδT cells (cluster 10) were overrepresented in livers of imq-treated mice and expressed *Mki67* gene (Fig. 5E).

**Figure 5:**
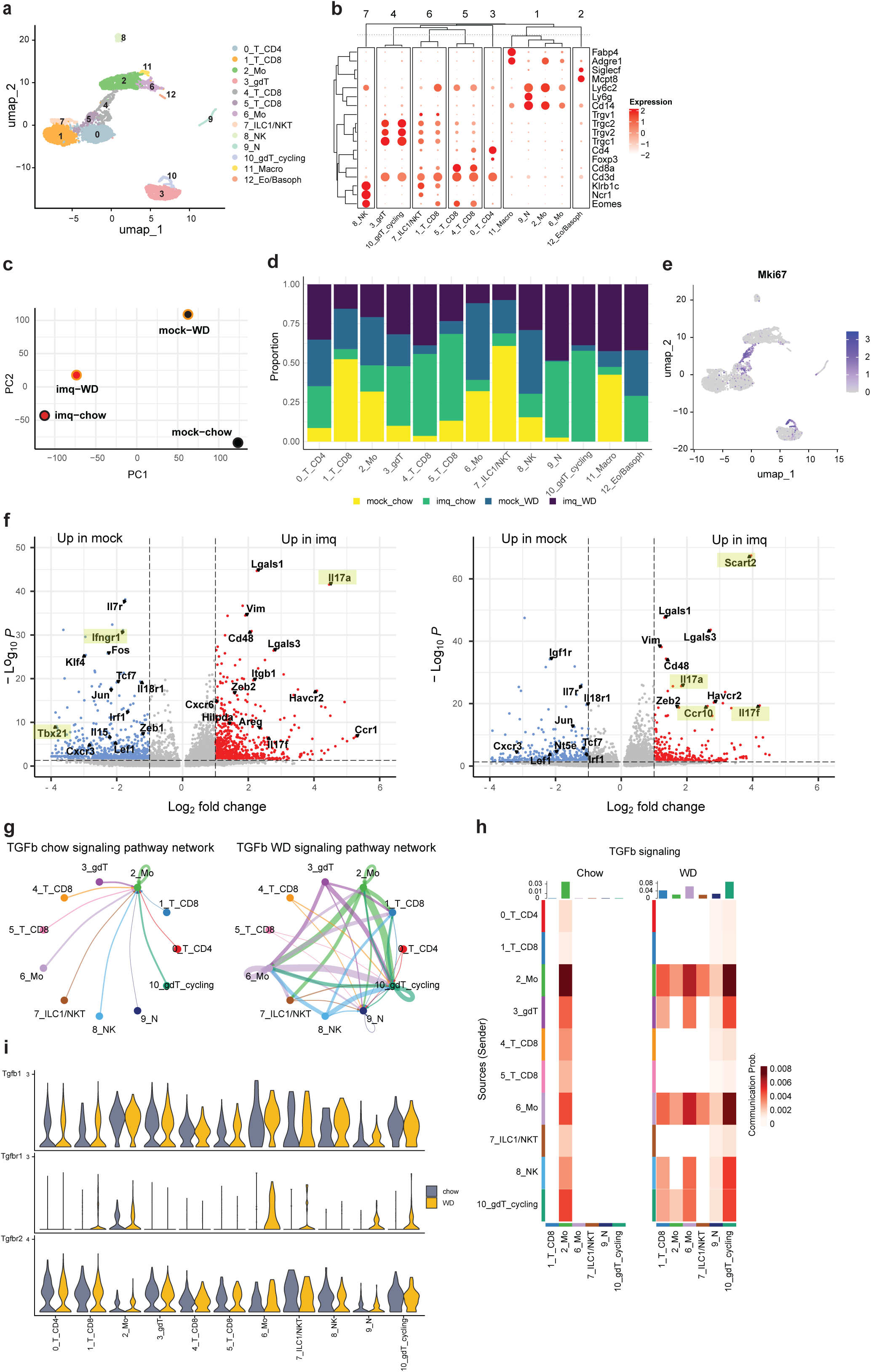
Single-cell RNA sequencing identifies hepatic γδT17 cells engaged in TGF-β signaling in early MASLD with concurrent chronic psoriasis. (**a**) UMAP projection of 7,118 liver immune cells, revealing 13 transcriptionally distinct clusters. (**b**) Hierarchical cluster annotation based on canonical lineage markers. (**c**) Principal component analysis (PCA) of all groups. (**d**) Proportions of identified clusters across groups. (**e**) Feature plot displaying *Mki67* expression across clusters. (**f**) Volcano plots of cluster 3 (γδT cells) comparing mock vs. imq treatment in chow-fed (left) and WD-fed (right) mice. (**g**) Circle plot depicting the TGF-β signaling network among clusters in imq chow (left) and imq WD (right) with line thickness indicating relative interaction strength. (**h**) Heatmap of TGF-β signaling among clusters in chow (left) and WD (right) groups. Rows represent sending clusters and columns receiving ones. Color intensity indicates communication probability, while bar plots summarize overall outgoing (top) and incoming (right) signaling strengths. (**i**) Violin plots showing expression of *Tgfb1*, *Tgfbr1*, and *Tgfbr2* across clusters, comparing chow (red) and WD (petrol blue). Each plot represents the distribution of normalized gene expression within the indicated cell cluster.

Non-cycling hepatic γδT cells (cluster 3) displayed significantly elevated *Il17a* expression in imq-exposed animals compared to mock-treated mice (Fig. 5F). In contrast, γδT cells from mock chow livers demonstrated higher levels of *Tbx21* and *Ifngr1*, characteristic of γδT1 cells. This signature was absent in mock WD, suggesting that WD may skew γδT cells toward a γδT17 phenotype. Supporting this notion, imq WD γδT cells expressed not only *Il17a*, but also *Il17f*, *Ccr10*, *Scart2*. As cluster 13 was not detectable in mock chow, comparisons were restricted to WD-fed mice, where *Il17a*, *Il17f*, and *Scart2* were significantly upregulated in imq WD relative to mock WD (Fig. S5B).

We next applied CellChat algorithm to explore intercellular communication between cell clusters. In imq WD condition, both γδT cell clusters, together with monocyte and NK cell clusters, exhibited increased interaction activity within the TGF-β signaling network, compared to imq chow condition, as reflected by a greater number and thickness of connecting lines between clusters in the CellChat circle plot. (Fig. 5G). Non-cycling γδT cells (cluster 3) acted as sender of TGF-β signals, with strong interaction detected in imq with monocytes/macrophages, proliferating γδT cells, and to a lesser extent with CD8^+^ Tcm and neutrophils (Fig. 5H).

Analysis of ligand expression revealed upregulation of *Tgfb1* in monocytes/macrophages (cluster 6) and NK cells (cluster 8) from imq WD livers (Fig. 5I), while γδT cells (clusters 3 and 10) expressed *Tgfb1* to similar levels across diets. Receptor analysis showed stable *Tgfbr2* expression across diets and clusters, whereas *Tgfbr1* was selectively upregulated in monocytes/macrophages (clusters 2 and 6), neutrophils (cluster 9) and cycling γδT cells (cluster 10). This is particularly relevant because canonical TGF-β signaling requires the presence of both Tgfbr1 and Tgfbr2 to form a functional receptor complex, meaning that only clusters expressing both receptors can be considered true functional targets of TGF-β signaling. Of note, *Col1a1*, a TGF-β target gene encoding type I collagen and an early marker of fibrosis, was significantly upregulated in livers of imq WD compared with mock WD (Fig. S5C).

Together, our data support that early MASLD and chronic psoriasis promote an IL-17 signature in hepatic γδT cells, which engage in TGF-β signaling and may contribute to MASLD pathogenesis through early fibrogenesis.

### Subclustering of hepatic γδT cells reveals a trajectory modulated by chronic psoriasis and MASLD in high-resolution

Given the significant role of *γδT cells in models of psoriasis-like dermatitis and MASLD*^41,42^, we next subclustered hepatic γδT cells and identified six transcriptionally distinct populations (Fig. 6A). Strikingly, clusters were largely segregated by treatment: clusters 0, 1, 4, and 5 were almost exclusively derived from imq-treated mice, whereas clusters 2 and 3 were restricted to mock-treated controls. Moreover, γδT cells from mock chow livers mapped only to cluster 2 and 3, while mock WD γδT cells were concentrated in cluster 2 (Fig. 6B). Based on gene expression, we annotated cluster 3 as Vγ1^+^ γδT1 cells (*Trgv1, Tbx21*, *Ifng*, *Cd27*), cluster 2 as Vγ6^+^ γδT17 cells (*Trgv6, Il1r1, Zbtb16*), and four clusters as Vγ4^+^ γδT17 cells: effector-like (cluster 0: *Klrk1, Rorc, Gzmb, Il17a/f)*, memory-like (cluster 1: high *Cd44, Il7r*), cycling (cluster 4: *Mki67*, *Ccnb1, Top2a*), and resident (cluster 5: *Itgae*, *Cd69*, high *Cd44*) (Fig. 6C). Whereas mock chow livers only contained Vγ6^+^ and Vγ1^+^γδT cells, mock WD livers also contributed to a minor fraction of Vγ4^+^ γδT cells (clusters 0, 1, and 4). Resident γδT17 cells were uniquely present in imq-treated mice (Fig. 6D). Pseudotime analysis constructs a cell trajectory among clusters representing a dynamic biological process. When this analysis was performed on our data, it suggested a developmental trajectory originating in effector-like Vγ4+ γδT17 cells (cluster 0), with potential to either enter the cell cycle (cluster 4), acquire a resident phenotype with heightened IL-23 sensitivity and Il17f expression capacity (cluster 5), or to memory-like cells (cluster 1), indicating more differentiated states, as shown by the lighter color gradient in the plot. (Fig. 6E). These results suggest that effector-like Vγ4⁺ γδT17 cells (cluster 0) may act as a central progenitor population, contributing to much of the γδT cell diversity observed in the liver. Together, our findings indicate that chronic psoriasis reshapes the composition, function, and developmental programming of γδT17 cells, which contribute to MASLD pathogenesis.

**Figure 6:**
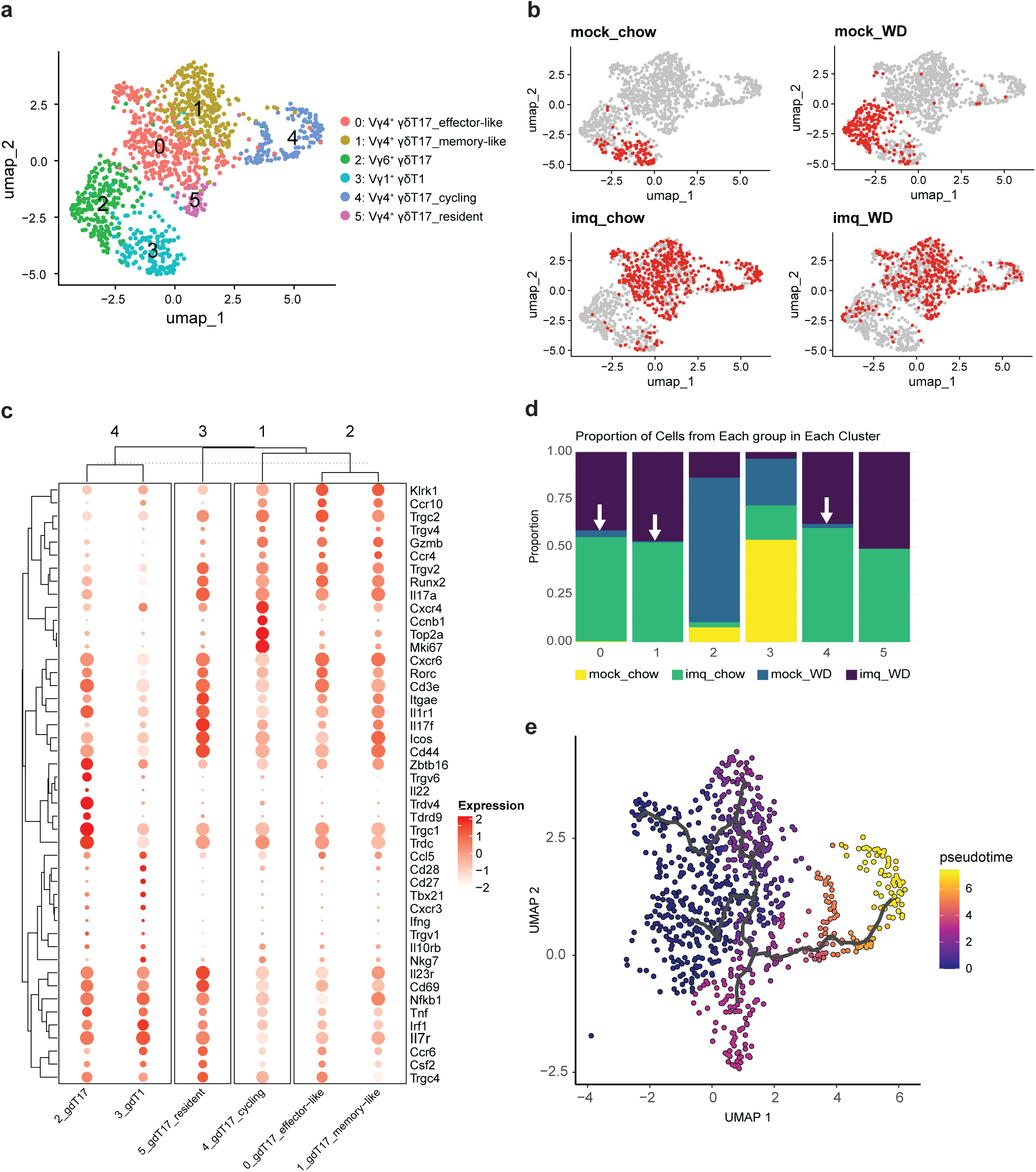
Subclustering of hepatic γδT cells reveals trajectories modulated by chronic psoriasis and MASLD. (**a**) UMAP projection of hepatic γδT cells. (**b**) UMAPs showing group-specific distribution of γδT cells (red) relative to all other groups (grey). (**c**) Hierarchical cluster annotation. (**d**) Proportions of identified γδT cell clusters across groups, orange arrows indicate presence of cells from mock WD livers. (**e**) Pseudotime analysis.

### Antipsoriatic IL-17 blockade improves hepatic steatosis in psoriasis patients with early-stage co-morbid MASLD

In a small prospective exploratory cohort of patients with moderate-to-severe psoriasis and co-morbid MASLD (n = 6, mean age 45.0 ± 5.4 years), treatment with clinically approved IL-17A/F inhibitor bimekizumab, administered according to standard dosing, resulted in a pronounced dermatological improvement (Fig. 7A). PASI scores declined significantly within 6 weeks of treatment (mean difference 9.5 ± 0.7; p < 0.0001; Fig. 7B), indicating a rapid and robust clinical response. Notably, hepatic steatosis, assessed by CAP, was significantly reduced after 6 months of therapy (327.2 ± 24.5 vs. 279.2 ± 18.2 dB/m; p = 0.024; Fig. 7C). At baseline, patients exhibited physiological liver stiffness (4.4 ± 0.4 kPa; reference < 6.0 kPa) and unremarkable non-invasive MASLD fibrosis scores, including FIB-4 (0.77 ± 0.07; reference < 1.3) and the NAFLD Fibrosis Score (NFS; −3.15 ± 0.53; reference < −1.46), which remained unchanged during treatment. However, we observed a trend toward a reduction in serum GGT (34.9 ± 6.6 vs. 26.5 ± 4.6 U/L; p = 0.062) and ALT levels (33.7 ± 8.8 vs. 25.1 ± 6.9 U/L; p = 0.067). No treatment-related adverse events were reported.

**Figure 7:**
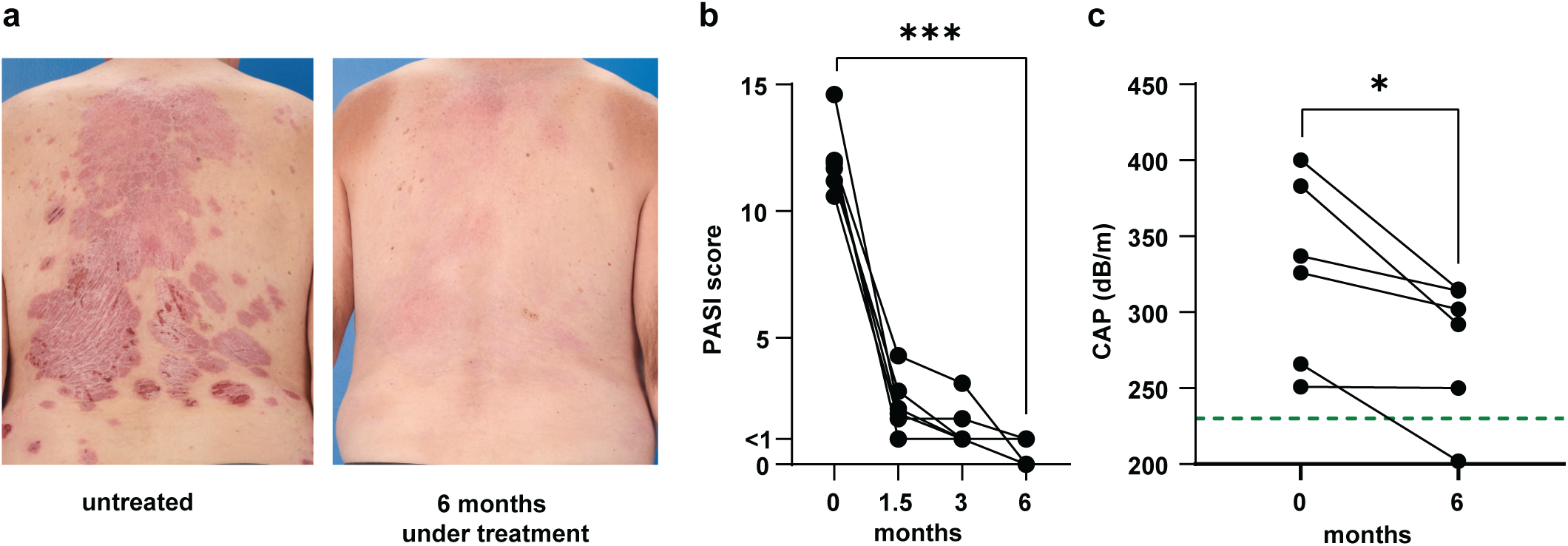
Clinical improvement of psoriasis and associated hepatic steatosis under IL- 17A/F inhibition. (**a**) Representative clinical photographs of a patient with moderate-to-severe plaque psoriasis before (left) and after 6 months of IL-17A/F-targeted therapy (right). (**b**) PASI scores during systemic treatment. (**c**) CAP, used as a surrogate for hepatic steatosis, before and after 6 months of therapy. Lines indicate individual patient. The dashed line represents CAP ≤ 230 dB/m, defining the absence of steatosis (S0). Statistical analysis was performed using paired t-tests.

## Discussion

Our study suggests that early MASLD and chronic psoriasis influence disease progression in a reciprocal manner, supporting the concept of inter-organ crosstalk between liver and skin. MASLD, induced by WD, was accompanied by increased skin immune cell infiltration and aggravated psoriatic skin inflammation upon imq treatment, while chronic relapsing psoriasis was linked to a shift of hepatic γδT cells towards an IL-17 phenotype, with Vγ4^+^ γδT participating in a TGF-β-associated signaling network that favors early fibrogenesis. Furthermore, MASLD and chronic psoriasis appeared to synergize in promoting systemic inflammation. How dietary components act on inflammatory responses in the skin has been studied extensively^22–27^. However, literature on mimicking MASLD-psoriasis comorbidity in mice is scarce and models often fail to recapitulate characteristics of human disease. Although psoriasis patients are at higher risk of developing MASH with progressive fibrosis^43^, the majority of patients have an asymptomatic steatosis that does not cause liver-related health issues^44^. Moreover, chronic plaque-type psoriasis is characterized by alternating periods of remissions and relapses, frequently triggered by stress, smoking, alcohol consumption, infections and drugs^45^. Therefore, our model was designed to mirror the typical clinical presentation of early MASLD, with cardiometabolic risk and hepatic steatosis in the absence of advanced inflammation or fibrosis, together with chronic psoriasis-like skin inflammation characterized by recurrent flares and remission phases. This model revealed that recurrent psoriatic dermatitis drives hepatic immunometabolic reprogramming via systemic expansion of Vγ4⁺ γδT17 cells, thereby promoting early liver remodeling. Importantly, consistent with observations in our novel murine model system, IL-17-targeted therapy improved hepatic steatosis in patients with moderate-to-severe psoriasis and co-morbid MASLD, which underscores the translational relevance of our study.

A previous report described more severe psoriatic inflammation due to comorbid MASH with fibrosis, although psoriasis itself did not impact MASH development in C57BL/6 mice^46^. However, the model relied on a single cycle of imq treatment combined with streptozotocin-induced β-cell destruction, resembling type 1 diabetes rather than the typical cardiometabolic context of MASLD, and included a most likely non-obesogenic 4-week period of HFD. In another study, one cycle of imq treatment combined with a short 3-week HFD supplemented with 7-ketocholesterol aggravated hepatic steatosis and exacerbated psoriasis-like dermatitis^24^. Although these findings support liver-skin inflammatory feedback driven by IL-17, immune cells have not been studied in detail. Based on literature^25^, we chose fructose-enriched WD, reflecting the human pathophysiology of MASLD more closely compared with HFD. In our study, 16 weeks of WD induced a phenotype consistent with early MASLD in patients, characterized by increased body and liver weight, as well as hepatic steatosis due to enhanced cholesterol and triglyceride storage, in line with upregulated *Mogat1*, *Cd36*, and *Scd1* expression. Induction of *Cxcl2* and *Ccl2* indicated recruitment of neutrophils and monocytes to the liver, while elevated plasma ALT levels indicated the onset of hepatocellular damage.

In line with the literature^22–25^, we observed a more pronounced skin inflammation upon imq treatment in WD-fed mice. Nevertheless, contrasting to previous reports^47–49^, rechallenge with imq did not lead to an aggravated phenotype. Although HFD and WD were shown to increase epidermal thickness upon acute imq treatment^22–25,46,50^, in our model, diet had no significant influence on imq-induced acanthosis. However, repetitive imq exposure likely drove epidermal thickening to a trophic limit, leading to desquamation, which was clinically observed in our model.

In line with previous reports^47,48^, we observed an increase of Vγ4⁺ γδT cells in both skin and sdLN upon imq treatment, displaying a γδT17 phenotype (CD44^hi^CD62L^lo^CD103^+^CD27^-^RORγt^+^). Vγ4⁺ γδT cells isolated from sdLN produced IL-17A, irrespective of diet.

Notably, the coexistence of early MASLD and psoriasis was associated with pronounced weight loss, increased hepatic immune cell counts, a pro-inflammatory plasma cytokine profile dominated by IL-17, and induction of hepatic acute-phase genes, including the IL-17 target *Lcn2* (Lipocalin-2)^51^. In addition, *Srebf1*, regulating hepatic lipid metabolism, was significantly upregulated. While SREBP-1c has been reported to directly repress *Il17a* transcription by competing with AHR at its promoter^52^, this regulation is unlikely to occur in our setting, as *Srebf1* is predominantly expressed in hepatocytes, whereas IL-17 is produced by immune cells. Instead, hepatic SREBP-1c drives lipogenesis, which may enhance oxysterol formation through increased cholesterol availability and oxidative stress. Oxysterols act as RORγt agonists^53^ and have been shown to promote γδT17-driven psoriatic skin inflammation^26^.

In psoriatic mice, hepatic upregulation of *Cxcl2* and *Ccl2* was paralleled by increased frequencies of neutrophils, monocytes/macrophages, and Vγ4⁺ γδT cells, as determined by flow cytometry. These Vγ4⁺ γδT cells could be subdivided into γδT1 and γδT17 subsets based on CD27 expression^54^, and their identity was further confirmed by intracellular staining of Eomes^55^ (γδT1) and RORγt (γδT17). While hepatic Vγ4⁺ γδT cells from mock-treated mice produced robust amounts of IFN-γ, those from imq-treated mice were highly enriched in γδT17 cells and lacked IFN-γ production. This functional skewing was further reflected by their failure to upregulate CD69 upon stimulation, despite elevated basal expression. This altered activation dynamic contrasts with sdLN-derived Vγ4⁺ γδT cells, suggesting a pre-activated state through prior TCR signaling or a resident/memory-like phenotype with reduced responsiveness. Besides Vγ4⁺ γδT cells, Vγ6⁺ γδT cells have the potential to produce IL-17 and can be distinguished based on *Scart2* and *Scart1* expression, respectively^56^. Vγ4⁺ γδT cells were reported to be the key IL-17 producing population in imq-treated skin^57^, although scRNA-seq analysis of skin revealed Vγ6⁺ γδT cells as an additional source^58^. Our scRNA-seq data of hepatic γδT cells indicates that *Scart2*-expressing Vγ4⁺ γδT cells prevail in the experimental model of early MASLD with chronic psoriasis. In our model, hepatic γδT cells of healthy mice were predominantly Vγ1⁺ with only minor contributions from Vγ6⁺ cells. In early MASLD, however, the proportion of *Il17a*-expressing Vγ6⁺ and Vγ4⁺ γδT cells increased, underscoring the role of IL-17 in MASLD development. Transcriptomic comparisons further revealed greater similarity among mice with early MASLD (fewer differentially expressed genes), whereas chow-fed controls (mock vs. imq) showed more pronounced differences. In the setting of comorbid psoriasis and MASLD, Vγ4⁺ γδT cells engaged in TGF-β-driven interactions with monocytes and macrophages, known to contribute to the induction of *Col1a1*, a marker of early fibrosis^59^. In the context of imq treatment, Vγ4⁺ γδT cells were shown to egress from draining lymph nodes in an FTY720-sensitive manner, and accumulated not only in inflamed, but also in non-inflamed skin^47^. However, it remains to be demonstrated whether these cells can also be detected in the liver.

Given the chronic nature of both psoriasis and MASLD, an important future question is whether γδT cells in our model actively traffic between skin and liver during the course of disease, expand locally after prior seeding, or derive from bone marrow-precursors that directly home to the inflamed organs. Previously, Vγ4⁺ γδT cells were reported to home to the dermis and dLN^60^. In a model of repetitive imq treatment, recruitment of circulating Vγ4⁺ γδT cells was shown to be dispensable for the memory response observed upon re-challenge^47^. In line with this, another study demonstrated that early during HFD feeding, Vγ4⁺ γδT cells accumulated primarily in sdLNs rather than in the skin itself, and only later appeared in higher numbers in peripheral blood and skin^23^. These findings suggest that systemic expansion of Vγ4⁺ γδT cells can occur under metabolic stress, whereas their contribution to local responses in psoriatic skin may be less dependent on recruitment.

Whether such dynamics also extend to the liver, where we observed IL-17–skewed γδT cells, remains to be elucidated. Future studies using fate-mapping or photoconversion strategies will be required to define γδT cell migratory behavior, and the contribution of adipokine-secreting adipose tissue to systemic immune activation warrants further investigation. Although murine and human γδT cell repertoires differ substantially, our data suggest conceptual and functional parallels in the pathogenic IL-17 axis in both psoriasis and MASLD.

In psoriasis patients, circulating Vγ9Vδ2 γδT cells have been reported to redistribute to inflamed skin^61^, suggesting analogous recirculation trajectories that could potentially extend to hepatic sites. Accordingly, investigating whether interference with γδT cell migration or activation, e.g. via S1P receptor modulation, attenuates systemic inflammation represents a promising translational direction. Notably, in our clinical cohort of moderate-to-severe psoriasis patients with co-morbid MASLD, IL-17-targeted systemic therapy improved hepatic steatosis, supporting the relevance of this immunological axis in humans. Moreover, our model is compatible with reporter and knockout systems across genetic backgrounds, providing a platform to explore γδT cell-directed or IL-17–modulating therapeutic strategies at the skin-liver interface.

Together, our findings support the concept of an immunological liver-skin axis in mice, mediated by γδT17 cells, and align with our clinical observations where IL-17-targeted therapy improved hepatic steatosis in patients with psoriasis and co-morbid MASLD.

## Methods

### Mice, diets, skin treatments, and scoring

Eight-week-old female C57BL6/J mice (Charles River) were fed either a Western-type diet (WD, 21% butter fat, 1.3% cholesterol, and 42 g fructose/L drinking water, similar to Charlton et al.^62^) or a low-fat control diet (chow) for 16 weeks. Mice were maintained under specific pathogen-free conditions on a standard light-dark cycle with *ad libitum* access to food and water. Skin treatments were administered in three cycles, each consisting of five consecutive days. Body weight was measured weekly and daily during treatment cycles. At weeks 9, 12, and 15, mice were anesthetized by intraperitoneal (i.p.) injection of ketamine (100 mg/kg) and medetomidine (0.2 mg/kg) in phosphate-buffered saline (PBS), followed by dorsal hair removal using an electrical shaver. Subsequently, 5%-imiquimod (imq) cream (Aldara^®^, Meda) or agent-free vaseline (mock) was applied (62.5 mg per treatment)^63^. Anesthesia was antagonized with i.p. atipamezole hydrochloride (1 mg/kg in PBS). On days 2-5 of each cycle, mice were anesthetized with isoflurane for skin treatment. At the end of each cycle, disease severity was recorded according to the modified Psoriasis Area and Severity Index (mPASI)^50,64^. One day after the third treatment cycle, mice were euthanized by CO_2_ inhalation. All animal experiments conform to the Directive 2010/63/EU and were approved by the local authorities (Lower Franconian Government, Würzburg, Germany, file number 55.2.2-2532-2-1475-31).

### RNA isolation, reverse transcription and quantitative PCR analysis

Total RNA was isolated from 30 mg liver tissue according to the manufacturer’s instructions (NucleoSpin RNA kit, Macherey-Nagel). From each sample, 1 µg RNA was reverse-transcribed (High-Capacity cDNA Reverse Transcription kit, Thermo Fisher Scientific). Quantitative real-time PCR (qPCR) was performed with 10 ng cDNA per reaction, using SYBR Green Master Mix (Thermo Fisher Scientific). Relative mRNA expression levels were determined by the comparative ΔΔCt method, normalized to the housekeeping gene Rplp0. Primer sequences are provided in supplementary table 1.

### Plasma cytokine concentrations and ALT activity

After sacrifice of the mice, cardiac puncture was performed to collect whole blood. The samples were allowed to clot for 30 minutes at room temperature, then centrifuged to separate plasma (10 min, 2,500 r.p.m). The supernatant was collected and aliquoted before storage at -80 °C. Plasma concentrations of IL-6, IL-10, IL-17A, IL17F, IL-22, TNF-α, and IFN-γ were quantified using a bead-based multiplex assay (LEGENDplex™ MU Th Cytokine Panel, *BioLegend*) according to the manufacturer’s instructions. Data were acquired by flow cytometry (CytoFLEX LX Flow Cytometer, Beckman Coulter), and analyzed using the LEGENDplex™ Data Analysis Software. Plasma alanine aminotransferase (ALT) was measured using a commercial kit (MAK052, Sigma-Aldrich). Absorbance was determined on a microplate reader (Infinite 200 PRO, Tecan).

### Liver lipid extraction

Liver cholesterol and triglycerides were extracted according to the Folch method^65^. Snap-frozen liver tissue was mechanically disrupted and homogenized (Ultra Turrax T8, IKA) in chloroform-methanol (1:2). Lipids were recovered from the chloroform phase, dried, resuspended in distilled water containing 5% NP-40, and vortexed vigorously. Total cholesterol and triglyceride concentrations were determined by enzymatic assays (CHOL2, Roche; Free Glycerol Reagent and Triglyceride Determination kit, Sigma-Aldrich, respectively) following the manufacturers’ instructions. Absorbance was read using a microplate reader (Synergy HT, BioTek).

### Histology and image analyses

Skin strips (approximately 2 x 0.4 mm) and half of the left lateral liver lobe were fixed in 2% paraformaldehyde (PFA) at 4°C overnight, incubated in 30% sucrose solution at 4°C overnight, rinsed with PBS, air-dried, and embedded in O.C.T. Compound (Sakura Finetek). Cryosections of 3 µm (skin) and 7 µm (liver) thickness were prepared and stored at -80°C. Skin sections were rehydrated through a graded ethanol series and stained with hematoxylin and eosin using an automated stainer in the routine service of the dermatopathology unit. Epidermal thickness was measured microscopically at three sides per section by a board-certified dermatopathologist blinded to the experimental groups. Cryoconserved liver sections were rehydrated for 5 min in PBS and rinsed with 60% isopropanol. Sections were stained for 15 min with freshly prepared Oil Red O working solution To prepare the working solution, 1 g of Oil Red O was dissolved in 200 mL of isopropanol; 160 mL of this stock solution was then mixed with 120 mL of distilled water, allowed to stand for 1 hour, and filtered before use. After staining, sections were rinsed with 60% isopropanol, washed in distilled water, counterstained for 30 s with hematoxylin (1:1 diluted with distilled water, Sigma-Aldrich), rinsed again with distilled water and mounted (Aquatex®, Sigma-Aldrich). For each mouse, the lipid-stained area was quantified in three high power fields using ImageJ software.

### Single-cell suspension preparation

Livers, spleens, skin-draining lymph nodes, and back skin were collected. After removal of subcutaneous adipose tissue, back skin was minced and digested in DMEM (Gibco) enriched with 1% FCS, 10 mM HEPES (pH 7.2–7.5, Gibco), 40 µg/mL DNase I, 2 mg/mL collagenase IV, and 2 mg/mL hyaluronidase (Sigma-Aldrich) for 45 min at 37°C and 100 r.p.m. The suspensions were vortexed and passed through 100-μm cell strainers. Livers were cut into small pieces and digested in DMEM containing 10 mM HEPES, 20 μg/ml DNase I, and 1 mg/ml Collagenase D (Roche) for 40 min at 37 °C and 100 r.p.m. Suspensions were filtered through 100-μm strainers and lymphocytes were enriched by 40%/80% Percoll (Cytiva) gradient centrifugation (860 g, 20 min, 21 °C, no brake). Cells from the interphase were washed with PBS and used for flow cytometry analysis and cell sorting. Spleens were dissociated through 100-μm strainers and subjected to red blood cell lysis (150 mM NH_4_Cl, 10 mM KHCO_3_, 0.1 mM EDTA, pH 7.2-7.4). Skin-draining lymph nodes were homogenized in 1.5 mL tubes using pestles and filtered through 100-μm strainers. All single cell suspensions were used for flow cytometry analysis or cell sorting.

### In vitro PMA/ionomycin activation

Single-cell suspensions from spleens, livers, and skin-draining lymph nodes were plated in a 96-well U-bottom plate and stimulated with 0.025 μg/mL phorbol myristate acetate (PMA, Sigma-Aldrich) and 1 μg/mL ionomycin (Sigma-Aldrich) in RPMI medium (Gibco) supplemented with 1% penicillin-streptomycin (Sigma-Aldrich), 10% FCS (Sigma-Aldrich), 50 μM ß-mercaptoethanol (Gibco). Cells were cultured for 3.5 h at 37°C, with 1 μg/mL brefeldin A (BFA, Sigma-Aldrich) added after 1.5 h.

### Flow cytometry and cell sorting

For flow cytometry analysis, dead cells were excluded by Zombie NIR™ and non-specific binding was blocked by anti-CD16/CD32 antibodies. Fluorochrome-conjugated monoclonal antibodies were purchased from BioLegend, BioXCell, Thermo Fisher Scientific, and BD Biosciences. Extracellular staining was performed for 30 min at 4°C. Intracellular staining was carried out overnight using the FoxP3/Transcription Factor Staining Buffer set. Cells were acquired on a Cytek Aurora flow cytometer using SpectroFlo software. For cell sorting, dead cells were excluded by 7-AAD staining, and non-specific binding was blocked by anti-CD16/CD32 antibodies. From liver single-cell suspensions, live CD45⁺Ly6G⁻CD19⁻ cells were gated. Within this population, CD90.2⁺CD3⁺ cells were identified and activated conventional T cells (CD44⁺γδTCR⁻) and γδT cells (γδTCR⁺) were sorted. From the CD90.2⁻CD3⁻ fraction, CD11b⁺ cells were sorted. Cell sorting was performed on a BD FACSAria III cell sorter.

### Single-cell library preparation and single-cell RNA sequencing

Group-wise pooled single-cell suspensions were multiplexed via TotalSeq™ anti-mouse hashtag antibodies (BioLegend) to enable pooling of different experimental groups for scRNA-seq, and labeled by anti-CD45 TotalSeq-A antibody for CITE-seq profiling following the manufacturer’s protocol. Four mice were used for each experimental condition. Briefly, cells were incubated with Fc receptor blocking reagent to minimize non-specific binding and then stained with unique hashtag oligonucleotide-conjugated antibodies for 30 minutes at 4°C. The pooled and antibody-labelled cells were processed using the 10x Genomics Chromium Single Cell 3′ v3.1 platform according to the manufacturer’s protocol. Following encapsulation and reverse transcription, cDNA was amplified, and feature barcode libraries for both hashtags and the protein-derived tag were generated in parallel with gene expression libraries. Libraries were quantified using a Bioanalyzer (Agilent) and sequenced on a P3 flow cell via an Illumina NextSeq 2000 platform targeting a sequencing depth of 50.000 reads per cell.

### scRNA analysis

From Illumina FASTQ files, a unique molecular identifier (UMI) count matrix was generated using Cell Ranger (10x Genomics). The resulting matrix was subsequently analyzed in R (v4.3.2) using the Seurat package (v5.1). Quality control filtering was applied to exclude cells with fewer than 1,000 detected UMIs and those with >5% mitochondrial UMI content. Data were normalized using the *LogNormalize* method and scaled with the *ScaleData* function. Cells were assigned to their corresponding experimental condition according to prior hashtagging information using *HTODemux*. Untagged cells and doublets were excluded from downstream analyses. Highly variable genes were identified and used for principal component analysis (PCA), followed by uniform manifold approximation and projection (UMAP) for dimensionality reduction, based on the first 20 principal components. Cell clusters were identified using the *FindNeighbors* and *FindClusters* functions, with the clustering resolution parameter set to 0.5. Differentially expressed genes (DEGs) were identified using *FindAllMarkers*, and cell type annotations were manually assigned based on canonical marker gene expression and reference databases. Trajectory analysis of scRNA-seq data was performed using the pseudotime algorithm implemented in the Monocle3 package (https://cole-trapnell-lab.github.io/monocle-release/). This algorithm constructs a single-cell trajectory representing a dynamic biological process, ordering cells from less differentiated to more differentiated states. The starting population was defined as *Cluster 0: γδT17 effector-like cells* within the re-clustered γδT cell compartment. Cells were ordered along the inferred trajectory based on similarities in their gene expression profiles, and pseudotime values were projected onto the UMAP embedding as a continuous color gradient representing progression along the trajectory. To infer cell–cell communication between distinct cell subsets, we employed the CellChat algorithm (http://www.cellchat.org/). CellChat quantitatively infers intercellular communication networks from scRNA-seq data based on ligand-receptor gene expression information, using a curated mouse signaling pathway database as reference. Communication probabilities for each ligand-receptor interaction were calculated directly from the Seurat object.

### Patient recruitment and clinical assessment

Adult patients with moderate-to-severe chronic plaque psoriasis (Psoriasis Area and Severity Index, PASI > 10) were prospectively enrolled prior to initiation of systemic therapy. Key exclusion criteria included previous systemic antipsoriatic treatment, hepatotoxic concomitant medication, and harmful alcohol consumption, as defined by a score ≤7 in the Alcohol Use Disorders Identification Test (AUDIT)^66^. The study was approved by the Medical Ethics Committee of the Julius-Maximilians-University of Würzburg (reference number 48/24-sc), and all participants provided written informed consent in accordance with the Declaration of Helsinki. Patients underwent transient elastography (TE) testing using FibroScan to assess controlled attenuation parameter (CAP) and liver stiffness as indicators of hepatic steatosis and fibrosis, respectively, at baseline and after 6 months of treatment. Systemic antipsoriatic therapy was administered according to German clinical guidelines using the IL-17A/F inhibitor Bimekizumab. Bimekizumab was administered according to the approved dosing regimen (320 mg s.c. at weeks 0, 4, 8, 12, 16, and then every 8 weeks). PASI scores were determined at baseline, 6 weeks, 3 months, and 6 months. Anonymized clinical images are shown with written informed consent.

### MASLD assessment

TE was performed to diagnose and classify MASLD^67^. Clinical diagnosis was based on an elevated CAP value and the presence of at least one cardiometabolic risk factor3. Harmful alcohol consumption was excluded using a standardized questionnaire. TE was supervised by a trained hepatologist blinded to participants’ medical history using a FibroScan Compact 530 device (Echosens), assessing CAP as a surrogate marker for hepatic steatosis and liver stiffness as a marker of fibrosis.

### Statistical analysis

Mice were randomly assigned to experimental groups. Flow cytometry data were analyzed with FlowJo software (BD Biosciences). Statistical analyses were performed using GraphPad Prism. Data normality was assessed with Shapiro-Wilk and Kolmogorov-Smirnov tests. Depending on data distribution, either analysis of variance (ANOVA) or the non-parametric Kruskal-Wallis test was applied. Comparisons between two independent groups were performed using a two-sided unpaired Student’s t-test for normally distributed data or the Mann–Whitney U test for non-normally distributed data. Longitudinal clinical parameters in psoriasis patients were compared using two-sided paired Student’s t-tests. Statistical significance was defined as follows: *p < 0.05, **p < 0.01, ***p < 0.001, ****p < 0.0001; n.s., not significant. In bar graphs, each symbol represents an individual mouse. Error bars indicate means ± SEM.

### Data availability

Single-cell RNA-seq data have been deposited at GEO and are publicly available as of the date of publication. This paper does not report original code. Any additional information required to reanalyze the data reported in this paper is available from the corresponding author upon request.

## Acknowledgments

This study was supported by the *Else Kröner-Fresenius-Stiftung* (TWINSIGHT-06 to JF) and the *Interdisziplinäres Zentrum für Klinische Forschung* (IZKF) Würzburg (IZKF-Bridging Z-3BC/16 to JF, AdvCS2 to AK). The authors gratefully acknowledge Elena Werle, Margit Ott, Claudia Rüth, Nadine Vornberger, Sofie Riedmann, Daniela Hefner, and Claudia Hart for their excellent technical assistance. We further thank Heike Hermanns, PhD, for providing access to laboratory equipment, and Muhammad Ashfaq Khan, PhD, for advice on the experimental diets. The authors are also grateful to the animal caretakers Kathrin Burghardt and Rainer Brandner. We sincerely thank Tomás Lapido for his creative and professional assistance in graphically realizing our data.

## Author contributions statement

A.K., J.F., G.G. and M.G. conceptualized the study. J.F. and Y.R. analyzed the data. J.F., T.O., L.F., and Y.R. performed the experiments. Y.R., R.D.L., and F.I. analyzed the scRNA-seq data. A.K. and F.K. performed histological scoring. A.G. provided hepatological expertise, supervised and interpreted transient elastography measurements. A.K., G.G., and M.G. provided intellectual input and supervised the experiments. A.K. and G.G. provided reagents, software, and equipment. J.F. drafted the manuscript.

---

**Supplementary Figure 1:**
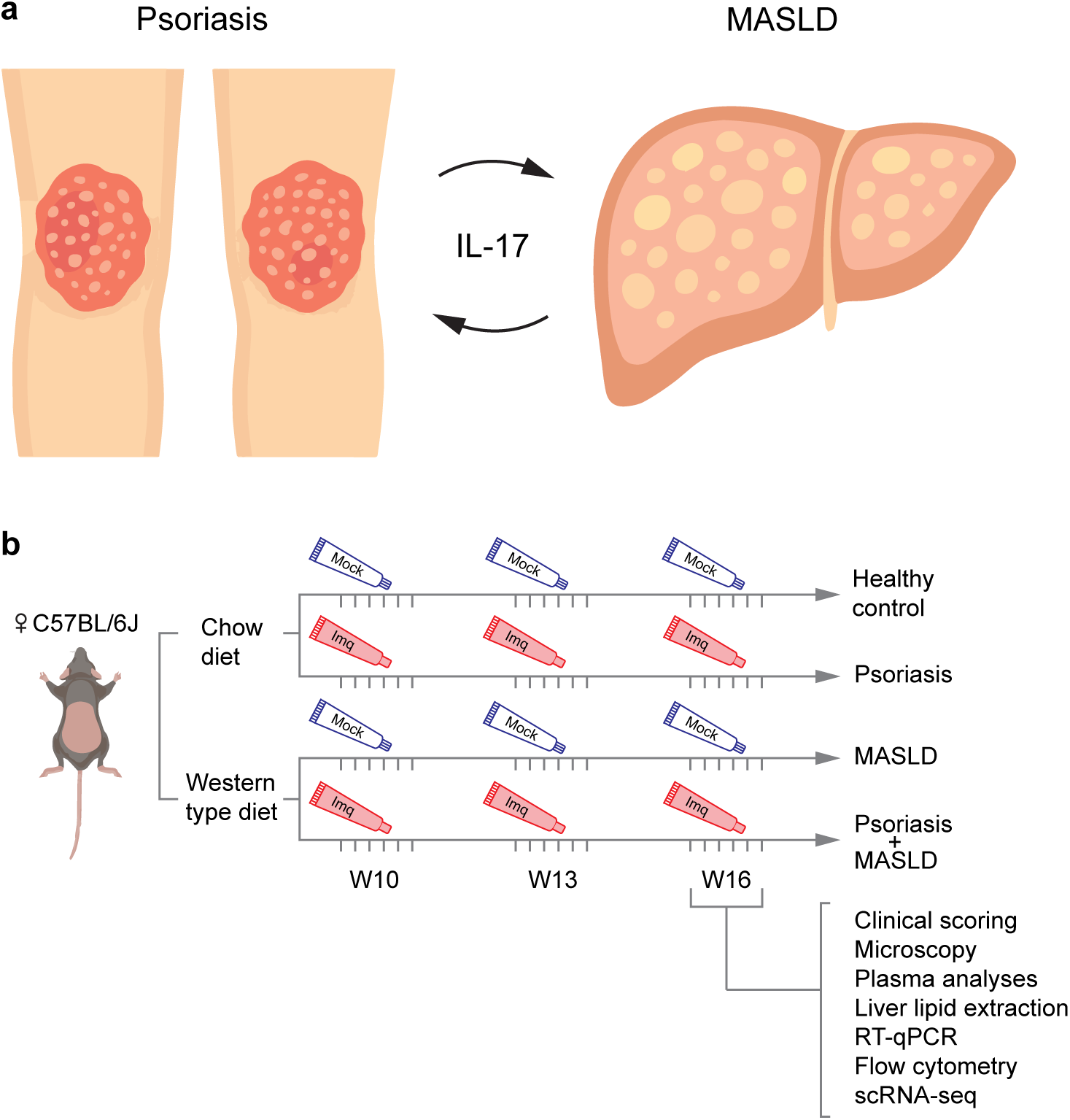
Hypothesis and experimental setup for modeling psoriasis and early MASLD in mice. **(a)** Psoriasis and metabolic dysfunction-associated steatotic liver disease (MASLD) co-occur in 50% of patients. Both diseases are driven by interleukin-17, suggesting a shared proinflammatory network. (**b**) Female C57BL/6J mice were fed a Western-type diet (WD) for 16 weeks to elicit early MASLD. To mimic chronic psoriasis-like skin inflammation, mice were topically treated by applying 5% imiquimod (imq) cream on the shaved back skin for 5 consecutive days, repeated three times. Controls were fed a normal chow diet and treated with Vaseline (mock). Four experimental groups were defined: healthy controls (mock chow), MASLD (mock WD), psoriasis (imq chow), and co-morbid psoriasis and MASLD (imq WD). Mice underwent clinical scoring, microscopy, plasma and liver lipid analyses, RT-qPCR, flow cytometry, and single-cell RNA sequencing (scRNA-seq).

**Supplementary Figure 2:**
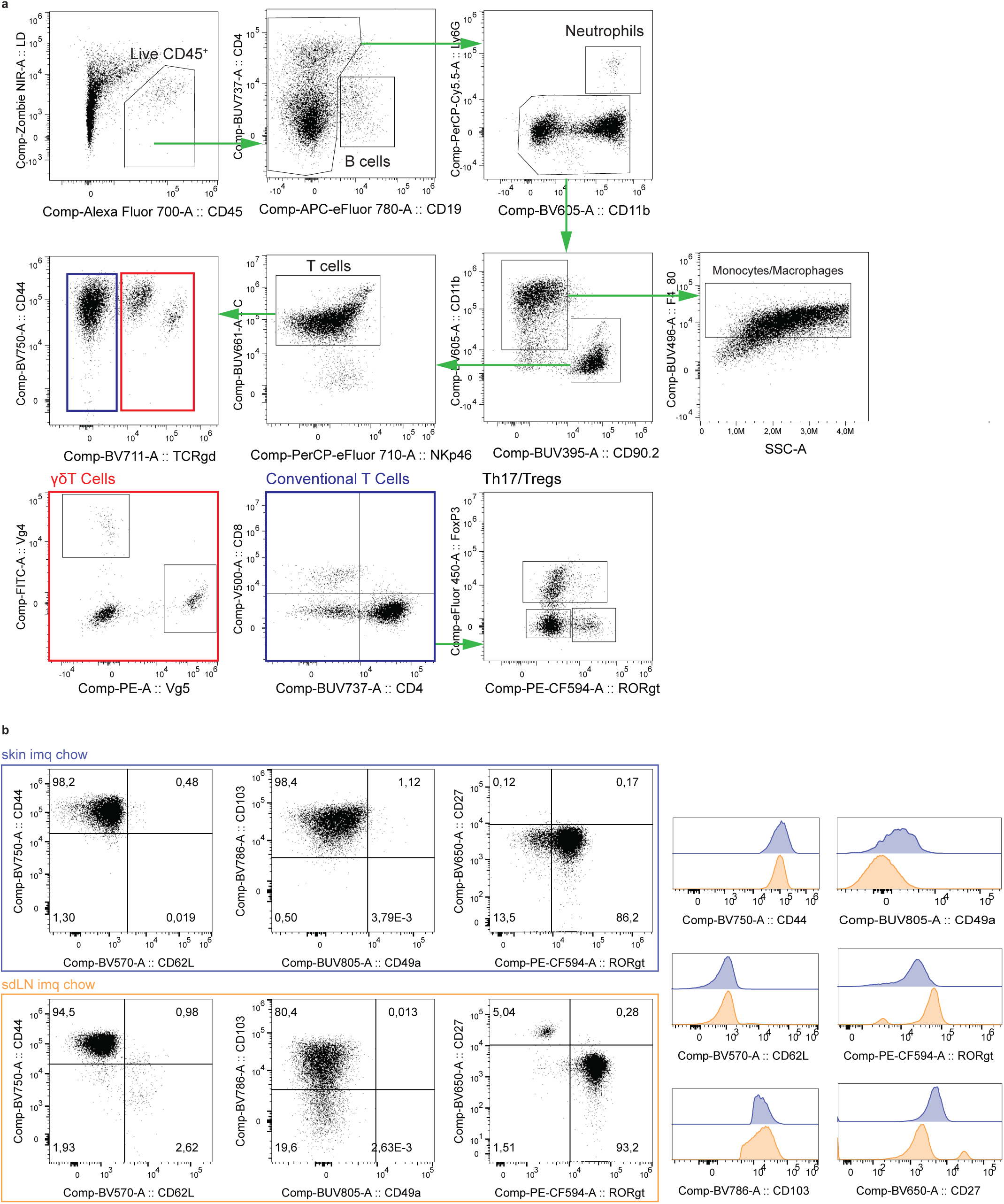
Gating strategy for skin immune cell subsets and phenotypic characterization of Vγ4⁺γδT cells in skin and draining lymph nodes. **(a)** Leukocytes were identified based on FSC/SSC properties and defined as live CD45⁺ cells after exclusion of singlets (not shown). B cells were defined as CD19⁺ cells. Neutrophils were identified as CD19⁻CD11b⁺Ly6G⁺, and monocytes/macrophages as CD19⁻Ly6G⁻CD11b⁺F4/80⁺ cells. T cells were gated as CD19⁻Ly6G⁻CD11b⁻CD90.2⁺CD3⁺NKp46⁻ cells and further divided into γδT cells (TCRγδ⁺) and conventional T cells (TCRγδ⁻). Within γδT cells, Vγ4⁺ and Vγ5⁺ subsets were distinguished; within conventional T cells, CD4⁺ and CD8⁺ populations were identified. Within CD4⁺ T cells, Th17 cells and Tregs were identified. Representative plots are shown (mock WD sample). (**b**) Representative flow cytometry plots (left) and histograms (right) showing expression of CD44, CD62L, CD103, CD49a, RORγt, and CD27 in Vγ4⁺ γδT cells isolated from skin and skin-draining lymph nodes (sdLN) of imq chow-treated mice. Histograms depict fluorescence intensity of the indicated markers in skin (purple) and sdLN (orange)

**Supplementary Figure 3:**
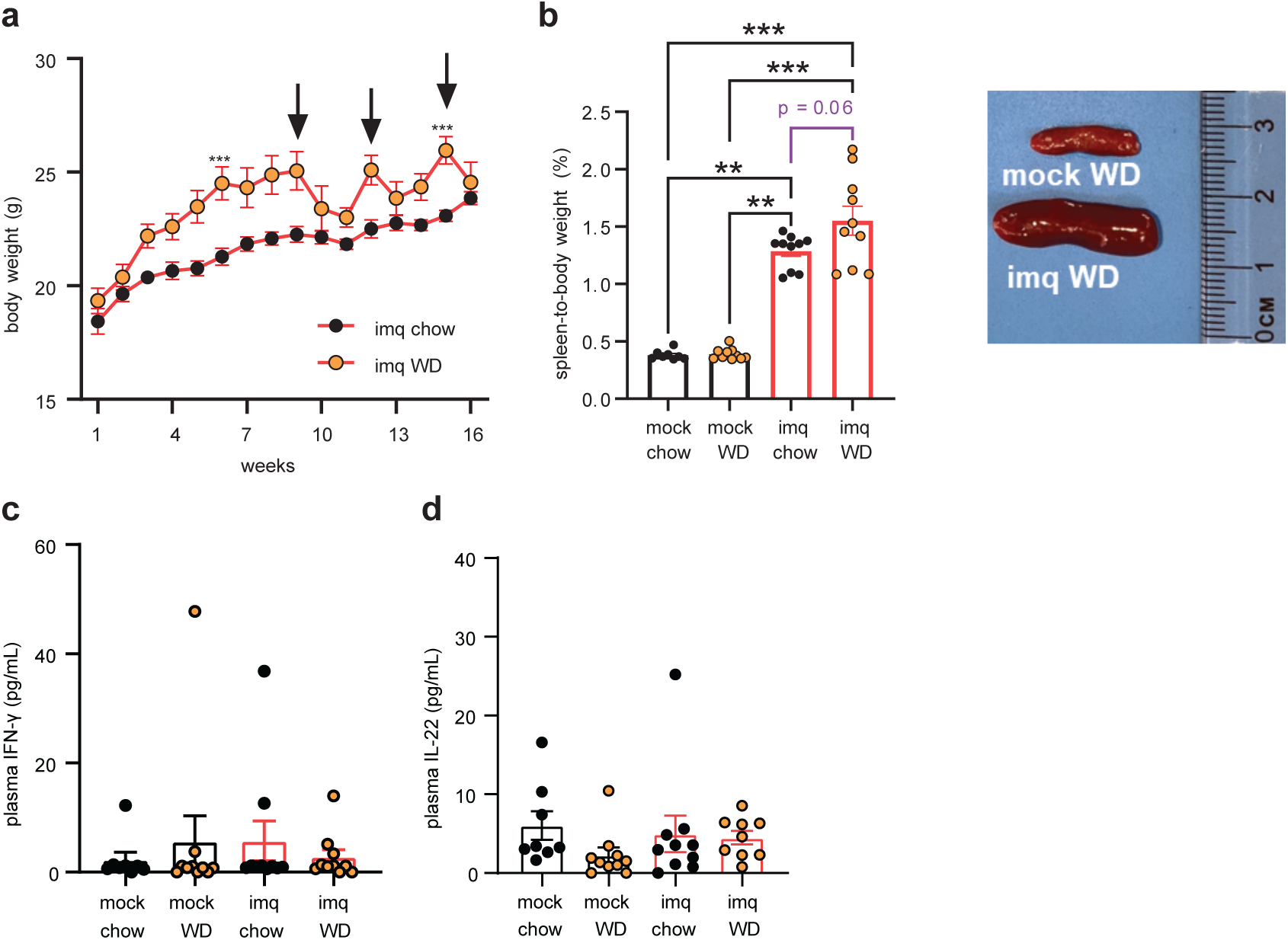
Body weight, spleen weight, and plasma cytokine levels. (**a**) Body weight (g) over the course of chow and WD. Data pooled from two independent experiments (n = 10 per group). Arrows indicate start of imq treatment cycles. (**b**) Spleen-to-body weight ratio (in percent, left) at sacrifice (mock chow, n = 8; imq chow, mock WD, and imq WD, n = 10 each) and representative photographs of spleens from mock WD and imq WD (right). (**c**) Plasma concentrations of IFN-γ and IL-22 (mock chow, n = 8; imq chow, mock WD, and imq WD, n = 10 each). Duplicate measurements per cytokine were averaged. Green dashed lines indicate mean concentrations in healthy control plasma as reported by the manufacturer. Kruskal-Wallis test, Mann-Whitney test and unpaired t-test (violet).

**Supplementary Figure 4.**
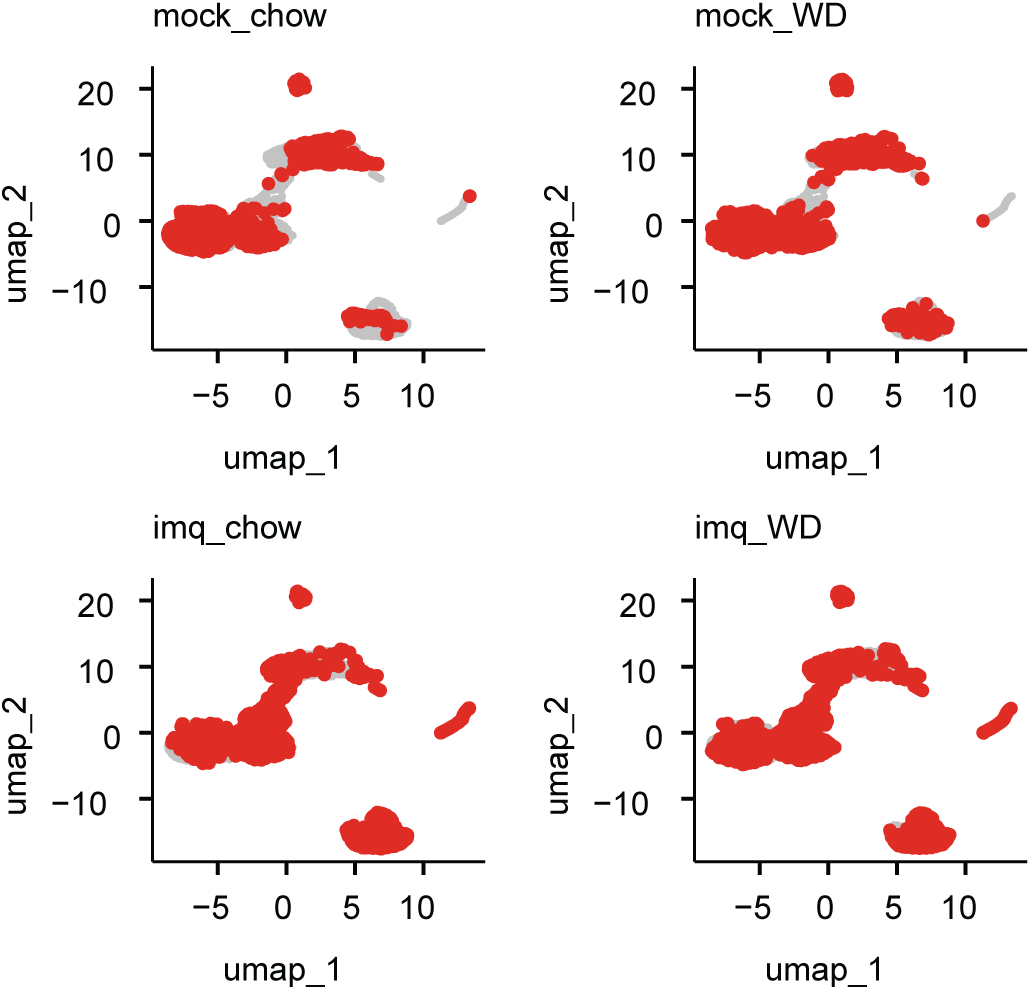
UMAPs showing group-specific distribution of cell clusters (red) relative to all other groups (grey). Each panel represents one experimental condition (mock chow, mock WD, imq chow, imq WD), visualizing the distinct clustering patterns depending on skin treatment and diet.

**Supplementary Figure 5:**
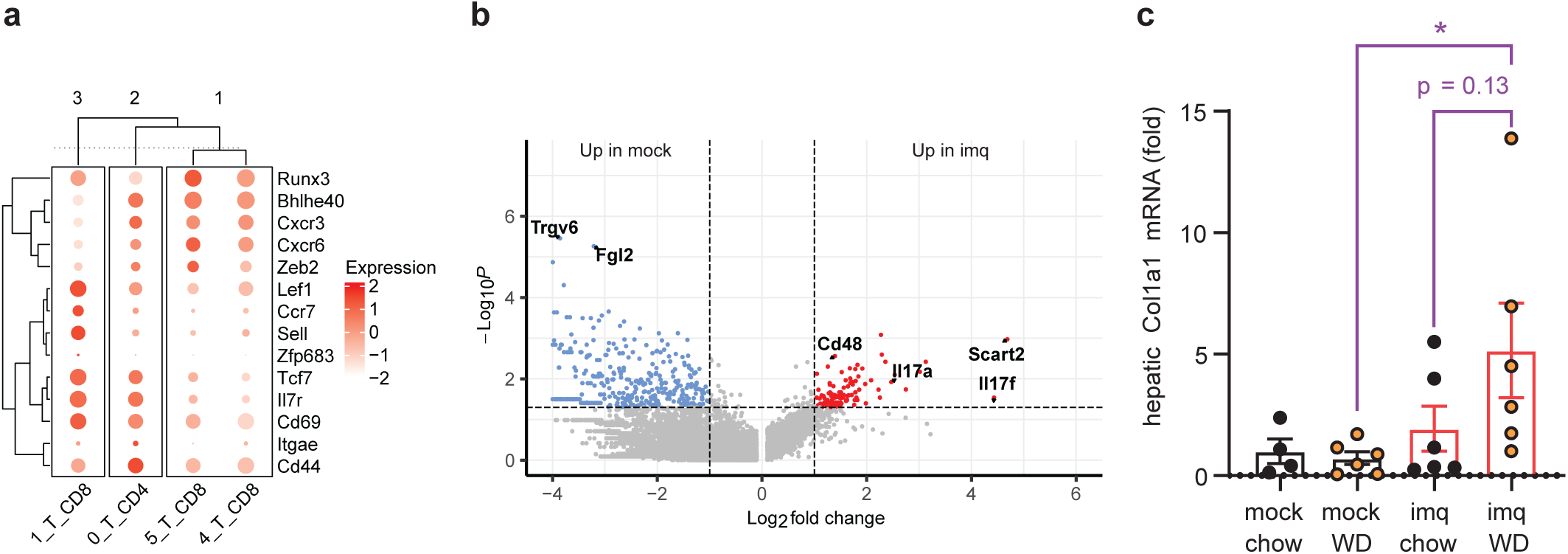
(**a**) Hierarchical dot plot showing expression of canonical markers used to annotate CD4⁺ and CD8⁺ T cell subsets. (**b**) Volcano plot of cluster 10 (cycling γδT cells) comparing mock vs. imq treatment in WD-fed mice. (**c**) Relative qPCR analysis of *Col1a1*. Results of mRNA expression were normalized to Rplp0 and presented relative to mock chow (mock chow, n = 4; imq chow, mock WD, and imq WD, n = 6 each). Mann-Whitney test (violet).

**Supplementary Table 1:**
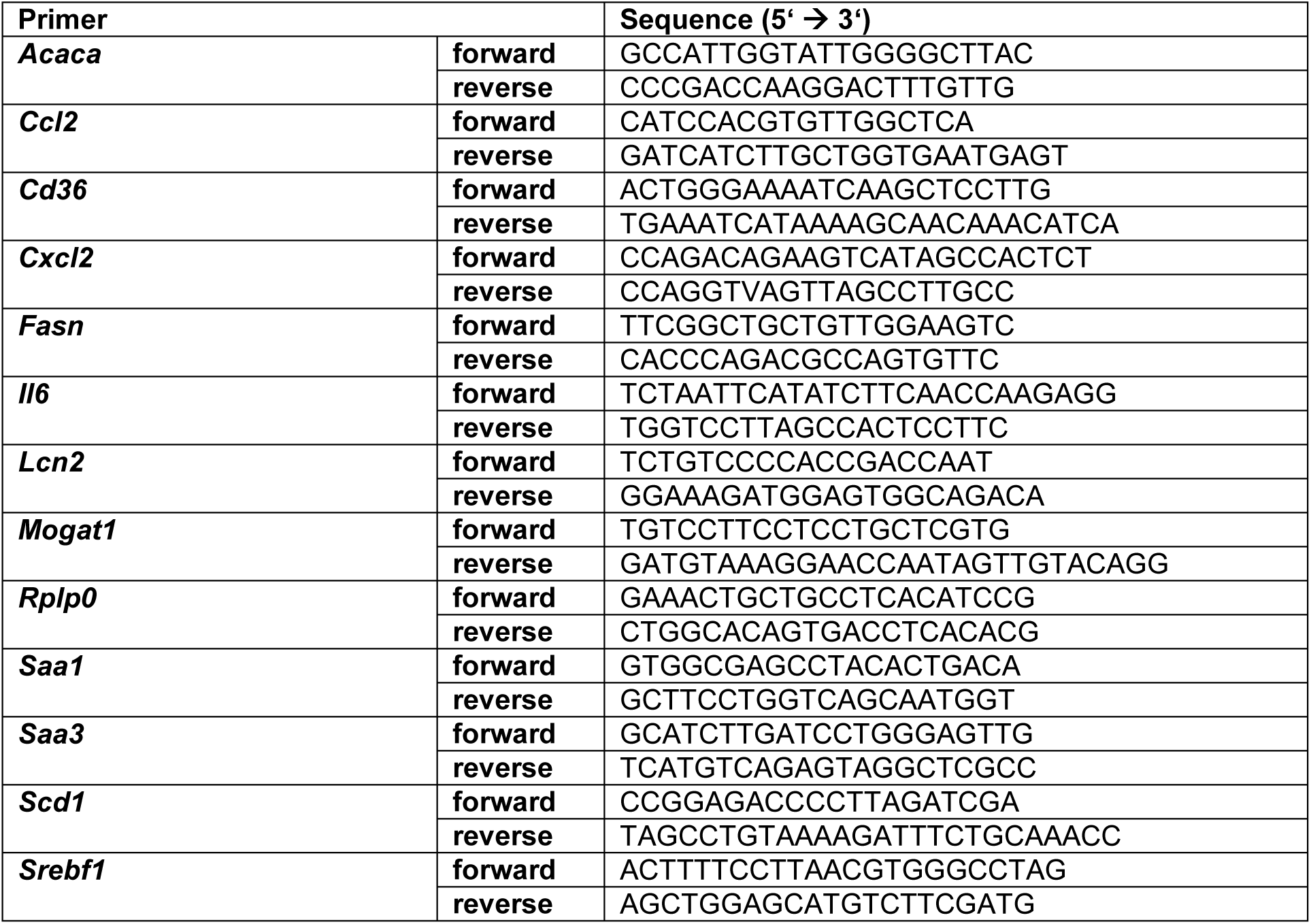
Primer pairs.

**Supplementary Table 2:**
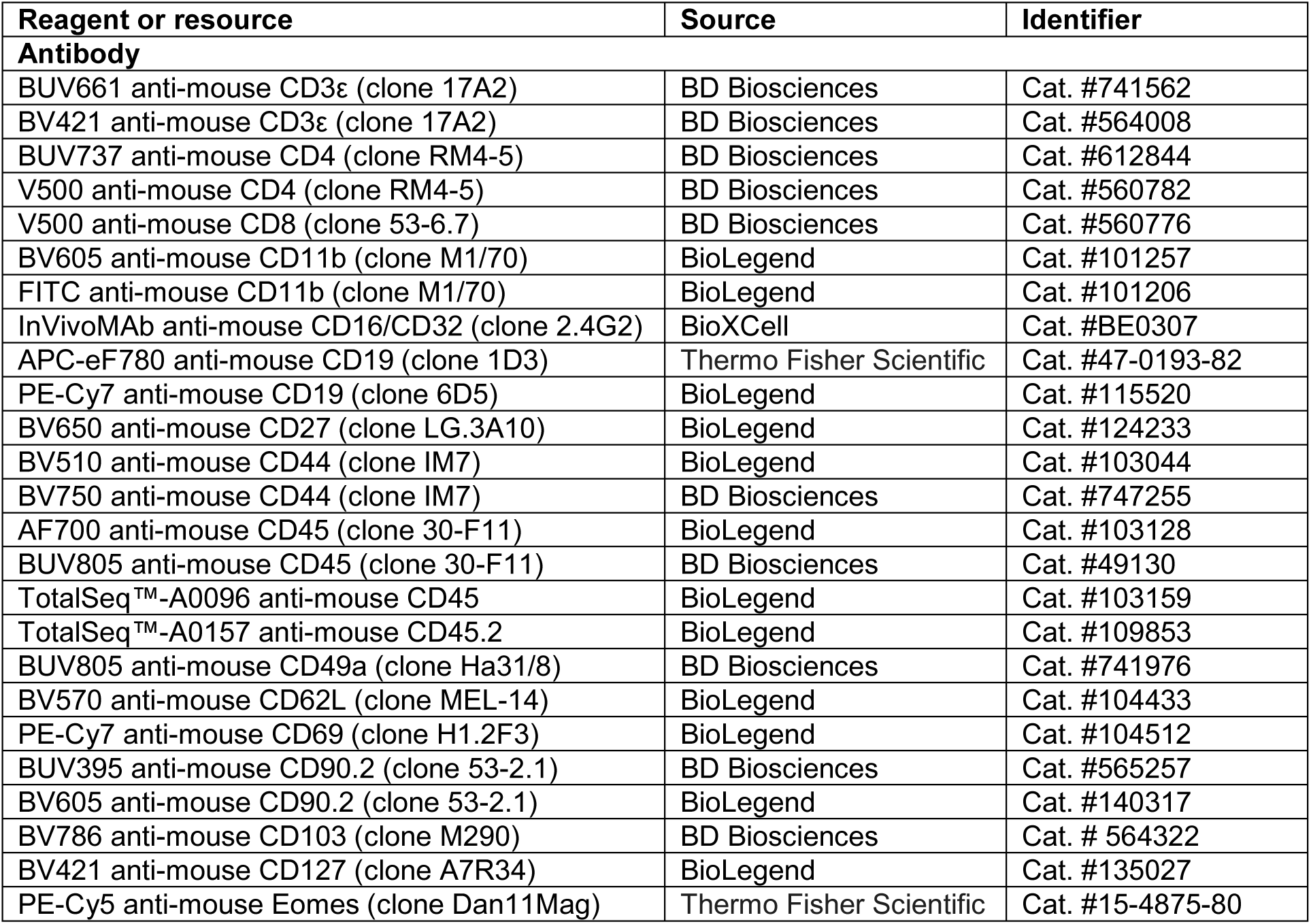
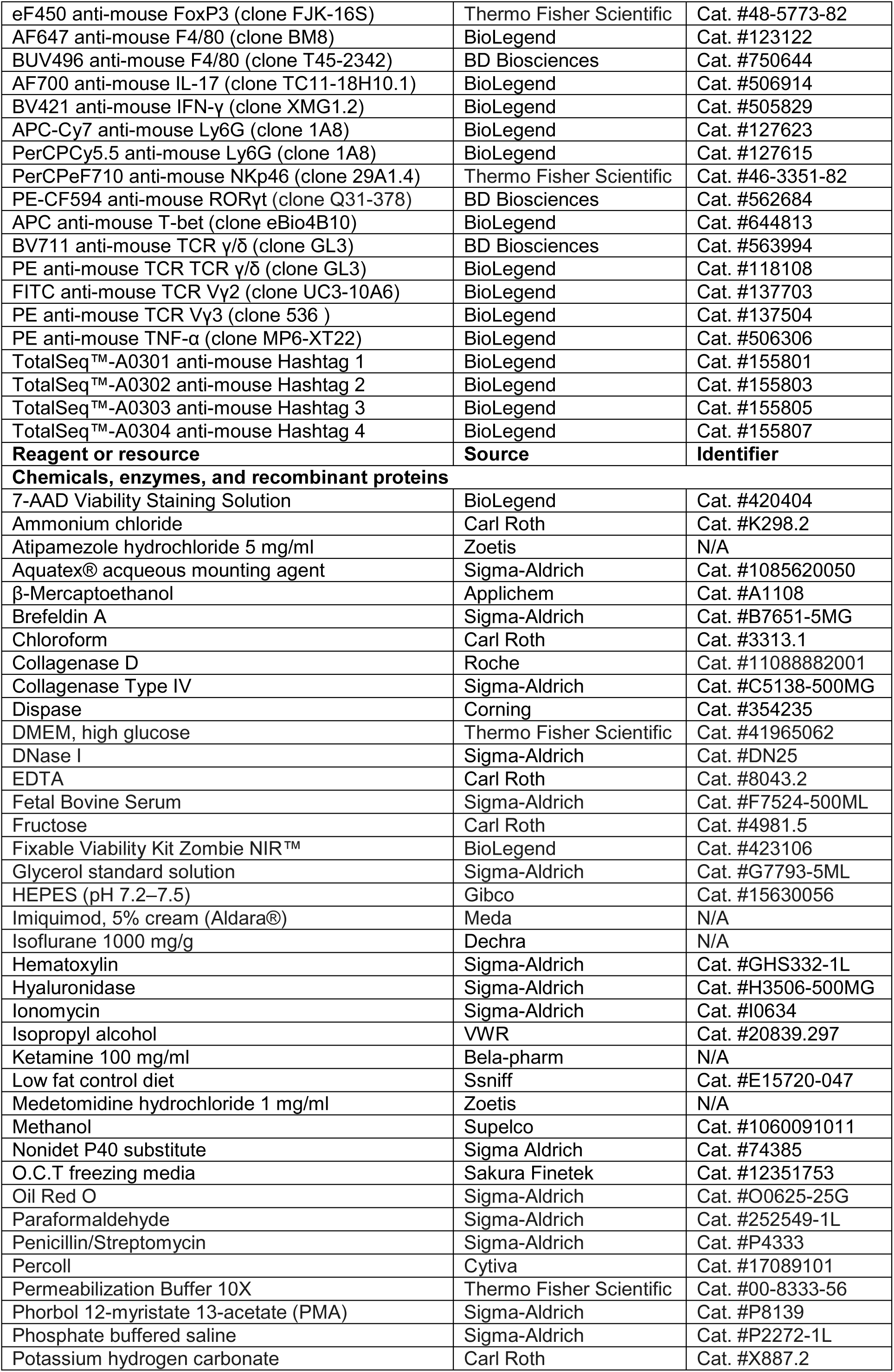
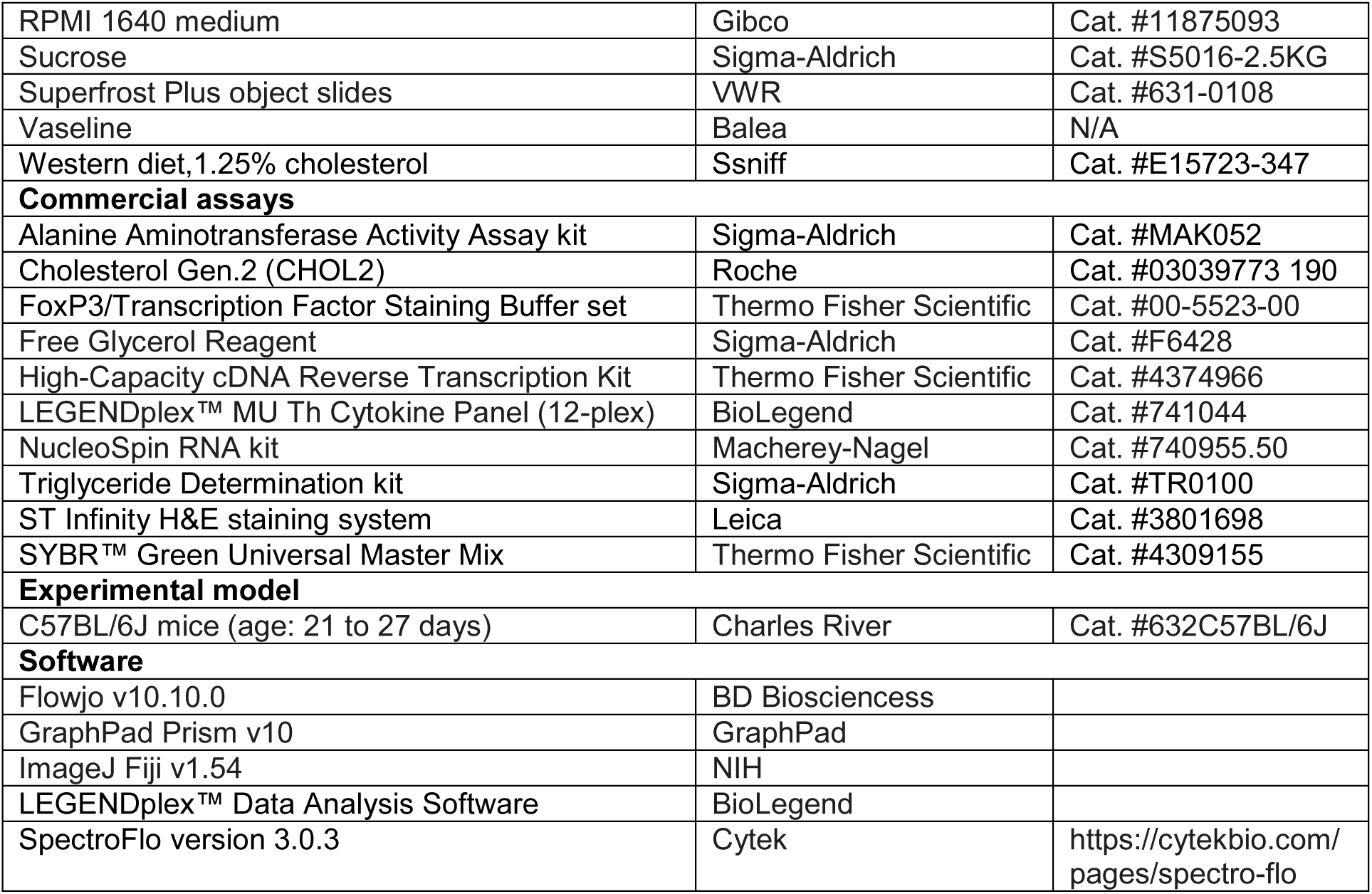
Materials and reagents.

| Reagent or resource | Source | Identifier |
| --- | --- | --- |
| **Antibody** |  |  |
| BUV661 anti-mouse CD3ε (clone 17A2) | BD Biosciences | Cat. #741562 |
| BV421 anti-mouse CD3ε (clone 17A2) | BD Biosciences | Cat. #564008 |
| BUV737 anti-mouse CD4 (clone RM4-5) | BD Biosciences | Cat. #612844 |
| V500 anti-mouse CD4 (clone RM4-5) | BD Biosciences | Cat. #560782 |
| V500 anti-mouse CD8 (clone 53-6.7) | BD Biosciences | Cat. #560776 |
| BV605 anti-mouse CD11b (clone M1/70) | BioLegend | Cat. #101257 |
| FITC anti-mouse CD11b (clone M1/70) | BioLegend | Cat. #101206 |
| InVivoMAb anti-mouse CD16/CD32 (clone 2.4G2) | BioXCell | Cat. #BE0307 |
| APC-eF780 anti-mouse CD19 (clone 1D3) | Thermo Fisher Scientific | Cat. #47-0193-82 |
| PE-Cy7 anti-mouse CD19 (clone 6D5) | BioLegend | Cat. #115520 |
| BV650 anti-mouse CD27 (clone LG.3A10) | BioLegend | Cat. #124233 |
| BV510 anti-mouse CD44 (clone IM7) | BioLegend | Cat. #103044 |
| BV750 anti-mouse CD44 (clone IM7) | BD Biosciences | Cat. #747255 |
| AF700 anti-mouse CD45 (clone 30-F11) | BioLegend | Cat. #103128 |
| BUV805 anti-mouse CD45 (clone 30-F11) | BD Biosciences | Cat. #49130 |
| TotalSeq™-A0096 anti-mouse CD45 | BioLegend | Cat. #103159 |
| TotalSeq™-A0157 anti-mouse CD45.2 | BioLegend | Cat. #109853 |
| BUV805 anti-mouse CD49a (clone Ha31/8) | BD Biosciences | Cat. #741976 |
| BV570 anti-mouse CD62L (clone MEL-14) | BioLegend | Cat. #104433 |
| PE-Cy7 anti-mouse CD69 (clone H1.2F3) | BioLegend | Cat. #104512 |
| BUV395 anti-mouse CD90.2 (clone 53-2.1) | BD Biosciences | Cat. #565257 |
| BV605 anti-mouse CD90.2 (clone 53-2.1) | BioLegend | Cat. #140317 |
| BV786 anti-mouse CD103 (clone M290) | BD Biosciences | Cat. # 564322 |
| BV421 anti-mouse CD127 (clone A7R34) | BioLegend | Cat. #135027 |
| PE-Cy5 anti-mouse Eomes (clone Dan11Mag) | Thermo Fisher Scientific | Cat. #15-4875-80 |
| eF450 anti-mouse FoxP3 (clone FJK-16S) | Thermo Fisher Scientific | Cat. #48-5773-82 |
| AF647 anti-mouse F4/80 (clone BM8) | BioLegend | Cat. #123122 |
| BUV496 anti-mouse F4/80 (clone T45-2342) | BD Biosciences | Cat. #750644 |
| AF700 anti-mouse IL-17 (clone TC11-18H10.1) | BioLegend | Cat. #506914 |
| BV421 anti-mouse IFN-γ (clone XMG1.2) | BioLegend | Cat. #505829 |
| APC-Cy7 anti-mouse Ly6G (clone 1A8) | BioLegend | Cat. #127623 |
| PerCPCy5.5 anti-mouse Ly6G (clone 1A8) | BioLegend | Cat. #127615 |
| PerCPeF710 anti-mouse NKp46 (clone 29A1.4) | Thermo Fisher Scientific | Cat. #46-3351-82 |
| PE-CF594 anti-mouse RORγt (clone Q31-378) | BD Biosciences | Cat. #562684 |
| APC anti-mouse T-bet (clone eBio4B10) | BioLegend | Cat. #644813 |
| BV711 anti-mouse TCR $\gamma/\delta$ (clone GL3) | BD Biosciences | Cat. #563994 |
| PE anti-mouse TCR TCR $\gamma/\delta$ (clone GL3) | BioLegend | Cat. #118108 |
| FITC anti-mouse TCR Vγ2 (clone UC3-10A6) | BioLegend | Cat. #137703 |
| PE anti-mouse TCR Vγ3 (clone 536 ) | BioLegend | Cat. #137504 |
| PE anti-mouse TNF-α (clone MP6-XT22) | BioLegend | Cat. #506306 |
| TotalSeq™-A0301 anti-mouse Hashtag 1 | BioLegend | Cat. #155801 |
| TotalSeq™-A0302 anti-mouse Hashtag 2 | BioLegend | Cat. #155803 |
| TotalSeq™-A0303 anti-mouse Hashtag 3 | BioLegend | Cat. #155805 |
| TotalSeq™-A0304 anti-mouse Hashtag 4 | BioLegend | Cat. #155807 |
| **Reagent or resource** | **Source** | **Identifier** |
| **Chemicals, enzymes, and recombinant proteins** |  |  |
| 7-AAD Viability Staining Solution | BioLegend | Cat. #420404 |
| Ammonium chloride | Carl Roth | Cat. #K298.2 |
| Atipamezole hydrochloride 5 mg/ml | Zoetis | N/A |
| Aquatex® aqueous mounting agent | Sigma-Aldrich | Cat. #1085620050 |
| β-Mercaptoethanol | Applichem | Cat. #A1108 |
| Brefeldin A | Sigma-Aldrich | Cat. #B7651-5MG |
| Chloroform | Carl Roth | Cat. #3313.1 |
| Collagenase D | Roche | Cat. #11088882001 |
| Collagenase Type IV | Sigma-Aldrich | Cat. #C5138-500MG |
| Dispase | Corning | Cat. #354235 |
| DMEM, high glucose | Thermo Fisher Scientific | Cat. #41965062 |
| DNase I | Sigma-Aldrich | Cat. #DN25 |
| EDTA | Carl Roth | Cat. #8043.2 |
| Fetal Bovine Serum | Sigma-Aldrich | Cat. #F7524-500ML |
| Fructose | Carl Roth | Cat. #4981.5 |
| Fixable Viability Kit Zombie NIR™ | BioLegend | Cat. #423106 |
| Glycerol standard solution | Sigma-Aldrich | Cat. #G7793-5ML |
| HEPES (pH 7.2–7.5) | Gibco | Cat. #15630056 |
| Imiquimod, 5% cream (Aldara®) | Meda | N/A |
| Isoflurane 1000 mg/g | Dechra | N/A |
| Hematoxylin | Sigma-Aldrich | Cat. #GHS332-1L |
| Hyaluronidase | Sigma-Aldrich | Cat. #H3506-500MG |
| Ionomycin | Sigma-Aldrich | Cat. #I0634 |
| Isopropyl alcohol | VWR | Cat. #20839.297 |
| Ketamine 100 mg/ml | Bela-pharm | N/A |
| Low fat control diet | Ssniff | Cat. #E15720-047 |
| Medetomidine hydrochloride 1 mg/ml | Zoetis | N/A |
| Methanol | Supelco | Cat. #1060091011 |
| Nonidet P40 substitute | Sigma Aldrich | Cat. #74385 |
| O.C.T freezing media | Sakura Finetek | Cat. #12351753 |
| Oil Red O | Sigma-Aldrich | Cat. #O0625-25G |
| Paraformaldehyde | Sigma-Aldrich | Cat. #252549-1L |
| Penicillin/Streptomycin | Sigma-Aldrich | Cat. #P4333 |
| Percoll | Cytiva | Cat. #17089101 |
| Permeabilization Buffer 10X | Thermo Fisher Scientific | Cat. #00-8333-56 |
| Phorbol 12-myristate 13-acetate (PMA) | Sigma-Aldrich | Cat. #P8139 |
| Phosphate buffered saline | Sigma-Aldrich | Cat. #P2272-1L |
| Potassium hydrogen carbonate | Carl Roth | Cat. #X887.2 |
| RPMI 1640 medium | Gibco | Cat. #11875093 |
| Sucrose | Sigma-Aldrich | Cat. #S5016-2.5KG |
| Superfrost Plus object slides | VWR | Cat. #631-0108 |
| Vaseline | Balea | N/A |
| Western diet, 1.25% cholesterol | Ssniff | Cat. #E15723-347 |
| **Commercial assays** |  |  |
| Alanine Aminotransferase Activity Assay kit | Sigma-Aldrich | Cat. #MAK052 |
| Cholesterol Gen.2 (CHOL2) | Roche | Cat. #03039773 190 |
| FoxP3/Transcription Factor Staining Buffer set | Thermo Fisher Scientific | Cat. #00-5523-00 |
| Free Glycerol Reagent | Sigma-Aldrich | Cat. #F6428 |
| High-Capacity cDNA Reverse Transcription Kit | Thermo Fisher Scientific | Cat. #4374966 |
| LEGENDplex™ MU Th Cytokine Panel (12-plex) | BioLegend | Cat. #741044 |
| NucleoSpin RNA kit | Macherey-Nagel | Cat. #740955.50 |
| Triglyceride Determination kit | Sigma-Aldrich | Cat. #TR0100 |
| ST Infinity H&E staining system | Leica | Cat. #3801698 |
| SYBR™ Green Universal Master Mix | Thermo Fisher Scientific | Cat. #4309155 |
| **Experimental model** |  |  |
| C57BL/6J mice (age: 21 to 27 days) | Charles River | Cat. #632C57BL/6J |
| **Software** |  |  |
| Flowjo v10.10.0 | BD Biosciences |  |
| GraphPad Prism v10 | GraphPad |  |
| ImageJ Fiji v1.54 | NIH |  |
| LEGENDplex™ Data Analysis Software | BioLegend |  |
| SpectroFlo version 3.0.3 | Cytek | https://cytekbio.com/pages/spectro-flo |

